# Chemoproteomic Profiling of PKA Substrates with Kinase-catalyzed Crosslinking and Immunoprecipitation (K-CLIP)

**DOI:** 10.1101/2025.03.23.644825

**Authors:** H. J. Bremer, M. K. H. Pflum

**Affiliations:** Wayne State University, Department of Chemistry, 5101 Cass Avenue, Detroit, MI 48202

**Keywords:** PKA, kinase, crosslinking, substrates, chemoproteomics

## Abstract

Phosphorylation is a highly regulated protein post-translational modification catalyzed by kinases. Kinases and phosphorylated proteins are key players in a myriad of cellular events, including cell signaling. When cell signaling networks are improperly regulated by kinases, various pathologies can arise, such as cancers and neurodegenerative disease. With critical roles in normal and disease biology, kinase-substrate interactions must be thoroughly characterized. Previously, the chemoproteomic method, kinase-catalyzed crosslinking and immunoprecipitation (K-CLIP), was developed to identify the kinases of a phosphoprotein substrate of interest. Here, K-CLIP was modified to profile the substrates of a kinase of interest. Specifically, the substrate profile of cAMP-dependent protein kinase (PKA) was studied with K-CLIP using a new ATP analog, ATP-alkyne aryl azide. Kinase-focused K-CLIP discovered SMC3 as a PKA substrate. With versatility for any kinase or phosphoprotein substrate of interest, K-CLIP will expand our understanding of kinase-mediated cell biology in healthy and diseased states.

## INTRODUCTION

Phosphorylation is a ubiquitous post-translational modification that regulates cell biology by controlling the enzymatic function, localization, and molecular interactions of cellular proteins. Kinases catalyze the transfer of the γ-phosphoryl from adenosine-5’-triphosphate (ATP) to a serine (Ser), threonine (Thr), or tyrosine (Tyr) residue of protein substrates (Figure 1A).^[1]^ Kinase-mediated phosphorylation is highly regulated and crucial for proper cell function. Irregular kinase activity is linked to a variety of pathologies, including cancer and neurodegenerative diseases, making kinases a prominent drug target.^[2]^ Monitoring kinase-mediated signaling events in normal and diseased cellular conditions is essential to understand cell biology and identify potential drug targets.

**Figure 1.**
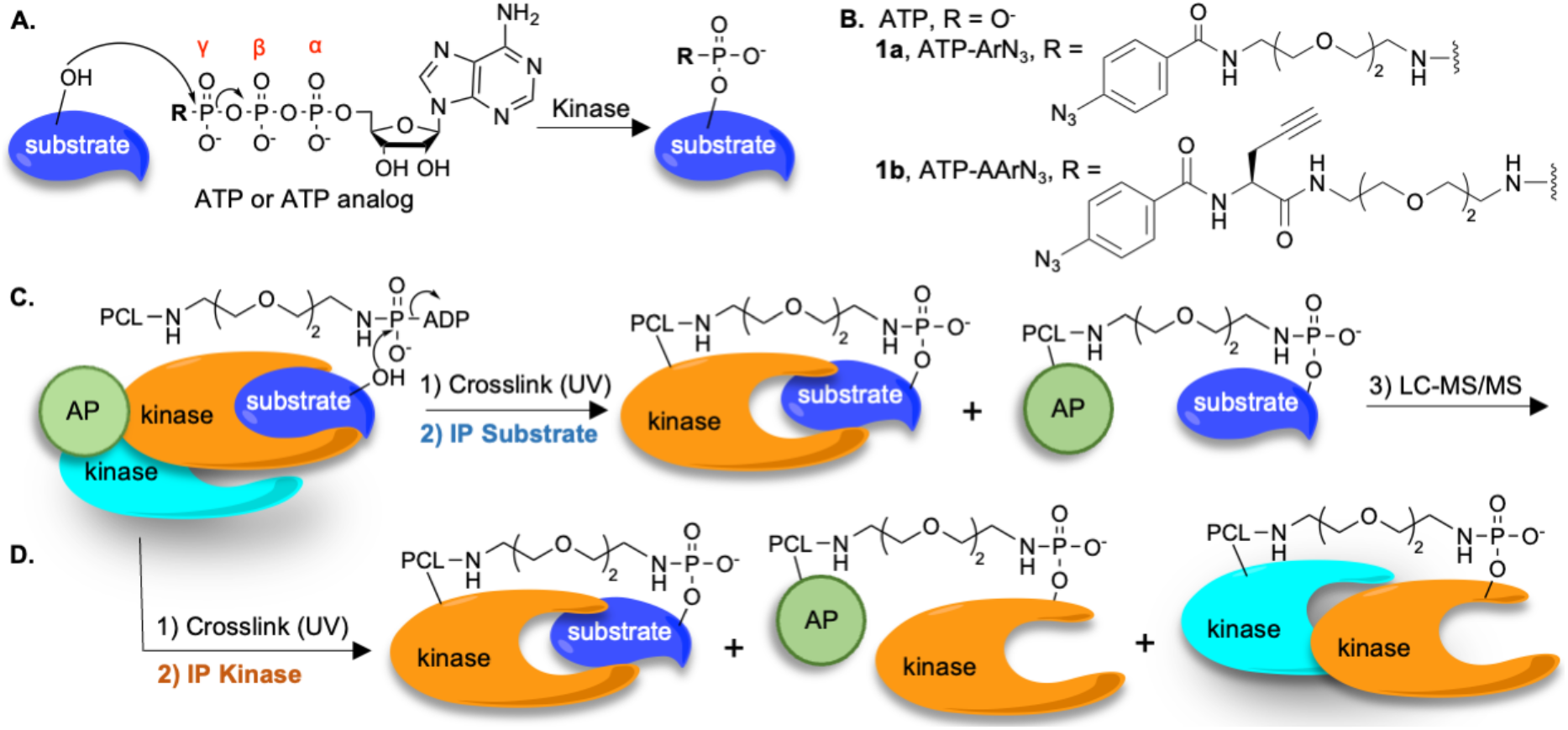
Kinase-catalyzed labeling and K-CLIP (A) Kinase-catalyzed phosphorylation mechanism, where the γ-phosphoryl of ATP or ATP analogs (shown in B) is transferred to a substrate protein. (B) Structure of ATP, ATP-aryl azide (ATP-ArN_3_), and ATP-alkyne aryl azide (ATP-AArN_3_). (C) In K-CLIP, kinase-catalyzed crosslinking of cellular lysates with ATP-ArN_3_ and UV light (step 1) will conjugate substrates to the kinase of interest (orange) via the photocrosslinker (PCL). Because the substrate can diffuse from the kinase before UV-activated crosslinking, substrates can also conjugate to nearby associated proteins (AP). Substrate-conjugated kinases and associated proteins are isolated by immunoprecipitation (IP) of the substrate (step 2) for identification using LC-MS/MS analysis (step 3). (D) After kinase-catalyzed crosslinking with ATP-ArN_3_ and UV (step 1), complexes isolated by immunoprecipitation of the kinase (orange, step 2) will include kinase-substrate complexes. In addition, because the kinase can be the substrate of another kinase (blue), upstream kinase-kinase or kinase-associated protein (AP) complexes will also be isolated, which complicates discovery of substrates by LC-MS/MS.

Due to the presence of more than 500 kinases and 20,000 phosphorylation sites, characterizing kinase-mediated signaling pathways remains a challenge.^[3]^ Mass spectrometry (MS) methods are currently used to efficiently identify phosphoproteins. However, MS methods alone fail to link phosphoprotein substrates to upstream kinases. To overcome the challenge of identifying kinase-substrate pairs, we previously introduced Kinase-catalyzed Crosslinking and ImmunoPrecipitation (K-CLIP) to identify the kinases of a target phosphoprotein substrate. K-CLIP relies on the fact that kinases accept γ-phosphoryl modified ATP analogs, such as ATP-aryl azide (ATP-ArN_3_, Figure 1B, **1a**), to label substrates with the γ-phosphoryl linked aryl azide crosslinker (Figure 1A). Coupled with UV irradiation to activate the aryl azide crosslinking group, the kinase and substrate will covalently conjugate in a phosphorylation dependent manner (Figure 1C, step 1).^[4-6]^ The resulting kinase-substrate complexes can then be enriched by immunoprecipitating the substrate of interest (Figure 1C, step 2). In addition to kinase-substrate complexes, K-CLIP also isolates substrate-associated protein complexes, which are generated when labeled substrates diffuse from the kinase before UV-activation of the aryl azide photocrosslinker. Analysis of the isolated complexes using liquid chromatography-tandem mass spectrometry (LC-MS/MS, Figure 1C, step 3) will identify both kinases and associated proteins of the phosphoprotein substrate, providing a powerful tool to probe both kinase-substrate and protein-protein interactions. Previously, K-CLIP identified both known and novel kinases of p53, as well as associated proteins.^[6]^

While K-CLIP successfully identified kinases of a chosen phosphoprotein substrate, discovering substrates of a selected kinase using K-CLIP is complicated because kinases can also be phosphorylated, acting as substrates to other kinases. In fact, kinases are often phosphorylated to initiate their phosphotransfer activity. For example, phosphoinositide-dependent protein kinase (PDK1) phosphorylates cAMP-dependent protein kinase (PKA).^[7]^ If PKA and PDK1 were conjugated using K-CLIP, distinguishing which protein acted as the kinase and which was the phosphoprotein substrate after LC-MS/MS analysis would be difficult. To further complicate kinase-focused K-CLIP, the kinase of interest after phosphorylation can diffuse away from the kinase complex before UV irradiation and crosslink to nearby associated proteins. In this case, conjugated associated proteins cannot be differentiated from substrates by LC/MS-MS analysis. In total, the challenge with kinase-focused K-CLIP is that upstream kinases, substrates, and associated protein will be co-enriched without a means of distinguishing substrates (Figure 1D).

Here, we developed a modified version of K-CLIP to overcome the challenge of distinguishing co-enriched substrates from associated proteins and upstream kinases of a target kinase. As an initial test of kinase-focused K-CLIP, substrates of PKA were discovered. PKA is an ideal model kinase, with well characterized enzymology and structure,^[8]^ as well as many known substrates.^[9]^ After kinase-focused K-CLIP and LC-MS/MS analysis, five hit proteins were identified, including two known PKA substrates, which documents the success of the method. Among the three novel hits, SMC3 was validated as a substrate of PKA. With the flexibility to study the kinase-substrate relationships of any target kinase or phosphoprotein substrate in any complex cellular mixture, K-CLIP is a powerful tool to discover kinase-substrate interactions in cell biology.

## RESULTS

### Design of kinase-focused K-CLIP

To develop a modified K-CLIP method for profiling substrates of a target kinase, a new bifunctional ATP analog was developed that allows enrichment of only phosphoprotein substrates, while avoiding upstream kinases or associated proteins. The new crosslinking analog, ATP-alkyne aryl azide (ATP-AArN_3_, Figure 1B, **1b**) is structurally similar to the original ATP-ArN_3_ **1a**, with a terminal aryl azide photocrosslinking group. However, ATP-AArN_3_ also contains an alkyne handle to enable secondary enrichment through click chemistry. In the modified kinase-focused K-CLIP strategy, cell lysates containing the kinase and substrates of interest are incubated with ATP-

AArN_3_ and UV irradiation for kinase-catalyzed crosslinking (Figure 2A, step 1). Immunoprecipitation of the selected kinase isolates all covalently crosslinked proteins, including substrates, upstream kinases, and associate proteins (Figure 2A, step 2). After elution from the antibody-bound beads, the kinase-containing complexes are attached to azide-modified agarose resin via click reaction with the alkyne of the ATP-AArN_3_ crosslinker (Figure 2A, step 3). Finally, the phosphoprotein substrates in each crosslinked complex are eluted using acid to selectively cleave the phosphoamidate bond within the linker (Figure 2B, step 4). The key feature of the second acid elution is selective release of only phosphoproteins from the bound beads, which will include only substrates and the original kinase of interest. Importantly, complicating upstream kinases and associated will remain on the resin. The selectively released kinase of interest and substrates are then collected for either gel-based or LC-MS/MS analysis. In total, the unique dual enrichment with ATP-AArN_3_ and kinase-focused K-CLIP has the potential to promote substrate profiling of any kinase of interest in any complex cellular mixture.

**Figure 2.**
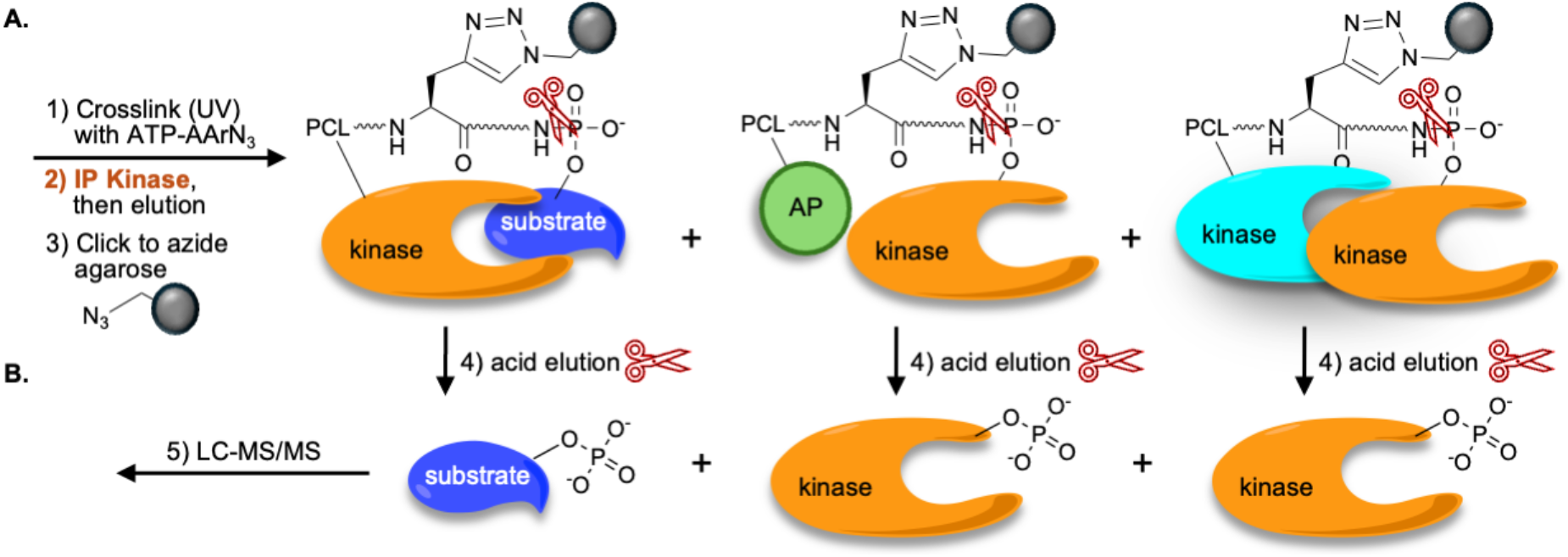
Kinase-focused K-CLIP (A) Kinase-catalyzed crosslinking of cellular lysates with ATP-AArN_*3*_ and UV light (step 1) will conjugate the kinase of interest (orange) via the photocrosslinker (PCL) to substrates, associated proteins (AP) and upstream kinases (blue), similar to Figure 1D. Kinase-conjugated proteins are isolated by immunoprecipitation (IP) and eluted (step 2), with subsequent attachment to azide-modified resin via click reaction of the alkyne group within the crosslinker (step 3). (B) Acid cleaves the phosphoramidate bond within the crosslinker (scissors, step 4), releasing the phosphoproteins from the protein-bound resin. The eluted phosphoproteins contain only the target kinase and substrates, which can be identified by LC-MS/MS (step 5).

### Synthesis of ATP-alkyne aryl azide

Synthesis of ATP-AArN_3_ first required preparation of Boc-protected polyethylene glycol (PEG) linker **2** (Scheme S1), as previously published.^[10]^ Fmoc-propargyl glycine **3** was coupled to linker **2** (Scheme 1). After Fmoc deprotection under basic conditions, the resulting amine intermediate of **4** was coupled to 4-azidobenzoic acid. Upon Boc deprotection under acidic conditions, the resulting amine intermediate of **5** was coupled to ATP to yield ATP-AArN_3_ **1b**, which was characterized by NMR and high-resolution mass spectrometry.

**Scheme 1.**
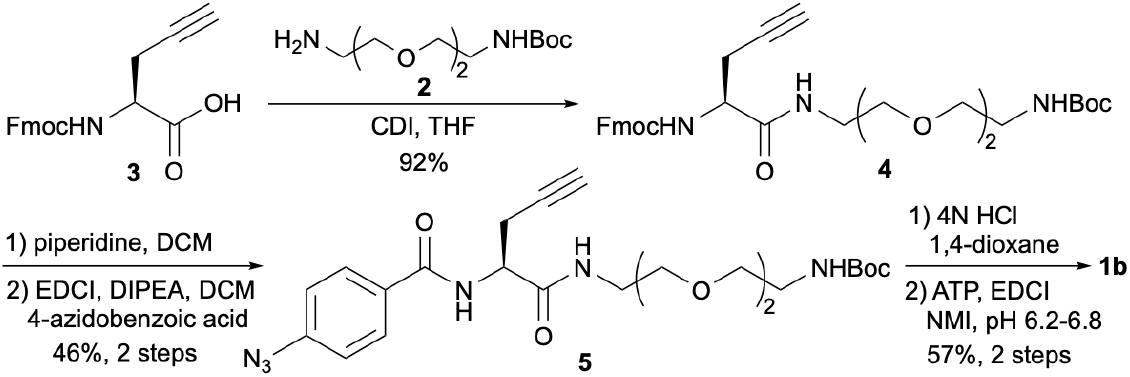
Synthesis of **1b**, ATP-AArN_3_

### Kinase-Catalyzed Crosslinking and K-CLIP with ATP-AArN_3_

Initial kinase-catalyzed crosslinking reactions were performed to ensure that ATP-AArN_3_ **1b** formed crosslinked complexes, similar to the original ATP-ArN_3_ **1a**.^[5, 11]^ ATP-AArN_3_ was incubated with and without UV light in a reaction with recombinant PKA kinase, which undergoes autophosphorylation to form PKA oligomer complexes after kinase-catalyzed crosslinking, as previously reported.^[12]^ Crosslinked complexes were separated via SDS-PAGE, electrotransferred to a membrane, and visualized via PKA western blot. PKA (43 kDa) and higher molecular weight PKA oligomers (86 kDa and 129 kDa) were observed with ATP-AArN_3_ and UV activation (Figure 3A, lane 3), but not without analog (Figure 3A, lane 1) or UV light (Figure 3A, lane 2). The PKA complexes are consistent with crosslinking observed with other photocrosslinker ATP analogs,^[5, 11, 12]^ which shows the crosslinking ability of ATP-AArN_3_.

**Figure 3.**
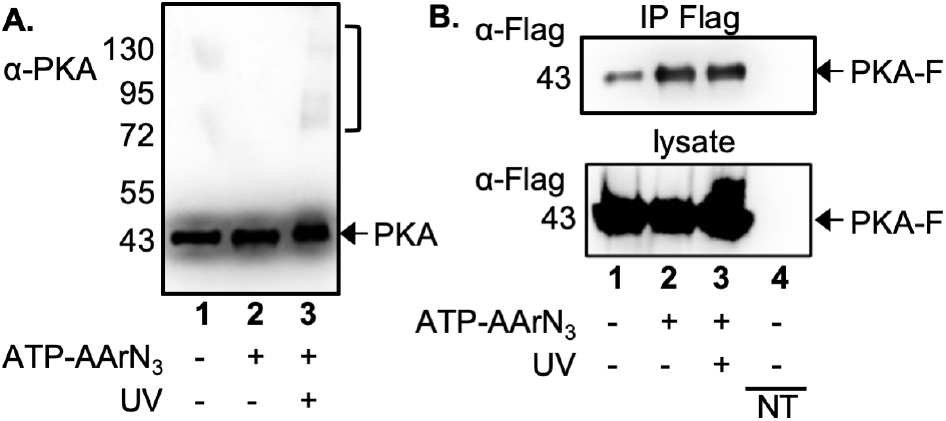
Kinase-catalyzed crosslinking and KCLIP with ATP-AArN_3_ and PKA. (A) PKA was incubated without (lane 1) or with ATP-AArN_3_ **1b** in the absence (lanes 1 and 2) or presence (lane 3) of UV for 2 hours at 31°C. After crosslinking, proteins were separated via SDS-PAGE and visualized with PKA (α-PKA) antibodies. The arrow indicates uncrosslinked PKA (43 kDa), and the bracket indicates crosslinked PKA dimers (86 kDa) and trimers (129 kDa). Additional independent trials are shown in Figure S11. (B) HEK293 cells were transfected without (lane 4, NT = nontransfected) or with a PKA-Flag expression plasmid (lanes 1-3). The resulting lysates were incubated with (lanes 2 and 3) or without (lanes 1 and 4) ATP-AArN_3_ (10 mM). PKA complexes were isolated via immunoprecipitation using the Flag tag. Enriched proteins were eluted using Flag peptide, and a secondary enrichment with agarose azide resin was performed using click chemistry. Proteins were eluted with acid, separated using SDS-PAGE, and visualized using Flag antibody (α-Flag). Additional independent trials are shown in Figure S12B. For both A and B, molecular markers are indicated to the left of the gel images.

As a next step to test ATP-AArN_3_, kinase-focused K-CLIP (Figure 2) was performed with PKA and analyzed by gel methods. PKA was overexpressed in HEK293 cells as a Flag tag fusion protein, and HEK293 cell lysates containing overexpressed PKA-Flag were incubated with ATP-AArN_3_ in the presence or absence of UV light for kinase-catalyzed crosslinking. Subsequent immunoprecipitation using Flag antibody-conjugated beads isolated all PKA crosslinked complexes (Figure 2A), which were then eluted using Flag peptide. We note that immunoprecipitation and elution via the Flag tag avoided heat and acid to maintain the phosphoramidate bond connecting the crosslinked proteins. After elution, click reaction with agarose azide resin isolated the PKA-bound complexes (Figure 2A), with PKA and its substrates selectively eluted under acidic conditions (Figure 2B). The expectation was the presence of PKA in crosslinked and uncrosslinked samples due to acidic release of phosphorylated PKA as both the kinase (Figure 2) and substrate (Figure S12A). After SDS-PAGE and western blot analysis, PKA-Flag was observed in crosslinked samples (Figure 3B, lane 3), consistent with successful enrichment with kinase-focused K-CLIP (Figure 2). PKA was also present in reactions without UV-mediated crosslinking (Figure 3B, lane 2) due to expected labeling and acidic release as a phosphoprotein substrate (Figure S12A). Reactions containing ATP-AArN_3_ isolated elevated levels of PKA (Figure 3B, lanes 2-3) compared to control reactions without ATP-AArN_3_ (Figure 3B, lane 1), where background signal was likely due to nonspecific binding to the resin. We note that nonspecific bead bound proteins will be accounted for in proteomic substrate discovery by including a negative bead control reaction. In total, gel analysis confirmed successful enrichment with ATP-AArN_3_ and kinase-focused K-CLIP, in preparation for substrate discovery.

### PKA K-CLIP for substrate discovery

Once established that kinase-focused K-CLIP successfully isolated PKA, proteomic analysis was used to identify bound PKA substrates. Enriched proteins after PKA-focused K-CLIP with HEK293 lysates containing overexpressed PKA-Flag were analyzed by LC-MS/MS with label-free quantitation using Proteome Discoverer.^[13]^ A total of 1618 proteins were identified in two independent trials (Figure 4A and Table S1A). As expected, PKAα (PRKACA) was present in both crosslinked (with UV) and uncrosslinked (without UV) samples, with lower intensity in the nontransfected control sample (Figure 4B, PRKACA), showing success of the method. In addition, although several PKA regulatory subunits were observed, none showed intensities similar to the catalytic subunits (Figure 4B), which indicated that the catalytic subunits were dissociated from the regulatory subunits and in an active form.

**Figure 4.**
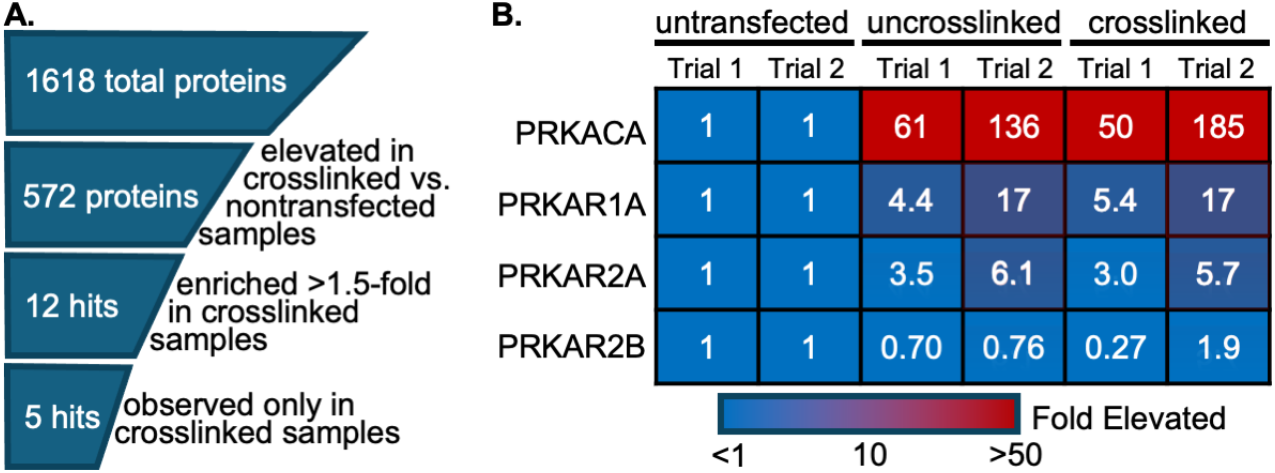
Analysis of PKA K-CLIP Proteomics data. (A) Enrichment analysis where the 1618 total proteins were initially evaluated to remove proteins enriched in nontransfected control samples compared to the crosslinked sample. Hit proteins were identified if enriched in crosslinked (with UV) versus uncrosslinked (without UV) samples by 1.5-fold. The top 5 hits were only observed in crosslinked, but not uncrosslinked samples. (B) PKAα catalytic subunit (PRKACA) and three PKA regulatory subunits (PRKAR1A, PRKAR2A, PRKAR2B) were observed in LC-MS/MS data and analyzed for enrichment in uncrosslinked or crosslinked samples by comparing to nontransfected samples (set to 1). The heat map color codes enrichment values with >50-fold enrichment in red and no enrichment (1-fold) in blue.

To identify possible PKA substrates, as an initial control, proteins with higher intensity in the bead binding negative control (without ATP-AArN_3_ or UV) compared to crosslinked (with ATP-AArN_3_ and UV) samples were removed from the hit list (Figure 4A). Hit proteins were then identified by comparing intensities of proteins in crosslinked samples (with ATP-AArN_3_ and UV) to intensities of the same proteins in uncrosslinked samples (with ATP-AArN_3_, but without UV). Using a threshold of 1.5-fold, 12 hit proteins were identified (Figure 4A, Table S1B). However, using a stricter cutoff with observable intensity in only crosslinked, but not uncrosslinked, samples, five candidate PKA substrates were selected for further consideration (Figure 4A, Table 1, Table S1B). Two of the final five hit proteins, serine/threonine-protein kinase NEK10 and cytidine triphosphate synthase 1 (CTPS1), are known substrates of PKA (Table 1),^[14, 15]^ demonstrating the success of K-CLIP for kinase substrate discovery. The three remaining proteins are known phosphoproteins, although without prior relationship to PKA.^[16-19]^

**Table 1.** PKA K-CLIP hit proteins.

| Protein name | Accession | Relation to PKA |
| --- | --- | --- |
| NEK10 | Q6ZWH5 | Known PKA substrate<sup>[14]</sup> |
| CTPS1 | P17812 | Known PKA substrate<sup>[15]</sup> |
| METTL7A | Q9H8H3 | Known phosphoprotein* |
| SEL1L3 | Q68CR1 | Known phosphoprotein* |
| SMC3 | Q9UQE7 | Known phosphoprotein* |

\*Documented phosphoprotein as reported in large scale phosphoproteomics and uniprot.org.<sup>[20]</sup>

### Validation of SMC3 as a PKA substrate

Among the three novel putative substrate hits identified using K-CLIP (Table 1), structural maintenance of chromosome protein 3 (SMC3) was selected for further validation. SMC3 is a known phosphoprotein with many phosphorylated serine and threonine residues,^[20]^ although no kinase has yet been identified for SMC3. To test if PKA phosphorylates SMC3, kinase assays were performed with γ-phosphate modified ATP analog, ATP-biotin (Figure S13A), in kinase-catalyzed biotinylation reactions (Figure 5A) using HEK293 lysates. For this cell-based assay, phosphorylation of cellular SMC3 by cellular PKA was monitored by the altering PKA kinase activity in HEK293 cells using the PKA inhibitor, H89, similar to prior studies.^[21]^ HEK293 cells were treated with H89 or vehicle control (DMSO), lysed, and incubated with ATP-biotin. Biotinylated phosphoproteins were isolated using avidin enrichment, eluted, separated via SDS-PAGE, electrotransferred to a membrane, and visualized using SMC3 specific antibody. SMC3 was biotinylated and enriched with ATP-biotin treatment (Figure 5B, lane 2) compared to the bead control without ATP-biotin (Figure 5B, lane 1), consistent with dynamic phosphorylation of SMC3 in HEK293 cells. Importantly, treatment of HEK293 cells with PKA selective inhibitor, H89, reduced the biotinylation and enrichment of SMC3 (Figure 5B, lane 3) compared to untreated samples (Figure 5B, lane 2), confirming that SMC3 biotinylation was PKA dependent. Quantified data from four independent trials showed that SMC3 biotinylation and enrichment is reduced with H89 treatment compared to untreated samples (Figure 5C). This cell-based assay confirmed that SMC3 is a substrate of PKA.

**Figure 5.**
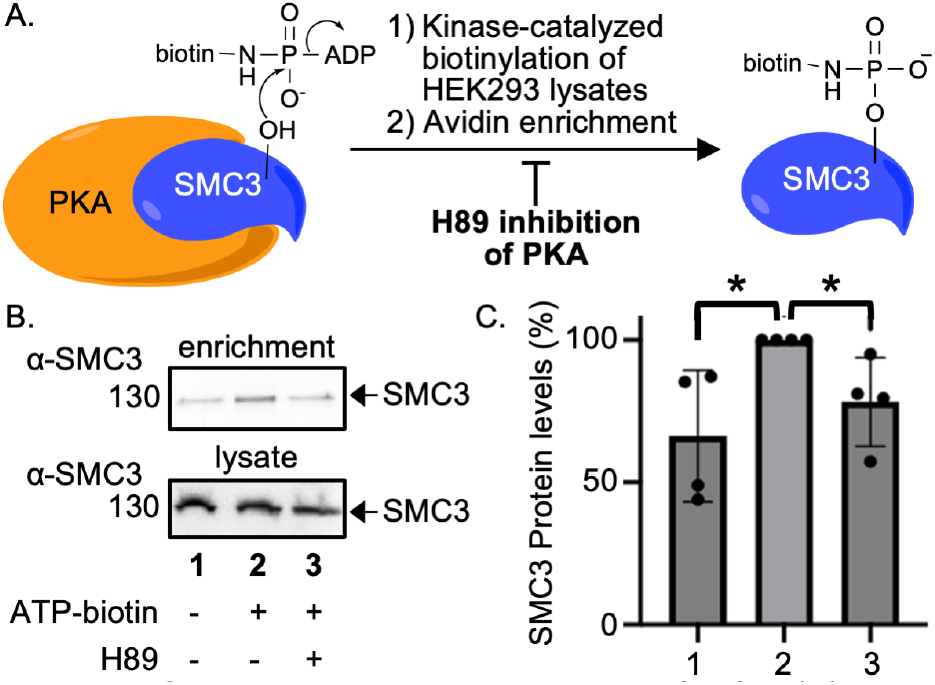
Cell-based assay with PKA and SMC3. (A) The cell-based assays involved incubation of untreated or H89 PKA inhibitor treated HEK293 lysates with ATP-biotin (Figure S13A), followed by avidin enrichment to isolate biotinylated SMC3, which was subsequently monitored using gel methods. (B) HEK293 cells were treated without (DMSO) or with PKA inhibitor H89 (30 μM). The resulting lysates were incubated with ATP-biotin (5 mM). Biotinylated proteins were enriched with NeutrAvidin resin, separated by SDS-PAGE, electrotransferred, and analyzed by immunoblotting with SMC3 antibody (α-SMC3). Input lysates before enrichment were separated as load controls. (C) Enriched SMC3 levels were quantified from four independent trials using ImageJ software and normalized as a percentage to lane 2 sample (set to 100%). Student t-test analysis was performed with GraphPad Prism (* p<0.05). Full gel images, independent trials, and quantification are documented in Figure S13.

## Discussion

Identifying the substrate profile of a kinase of interest has remained a challenge due to the difficulty of isolating the transient kinase-substrate complex. To remedy this obstacle, photocrosslinking ATP analogs were previously created to covalently link a kinase to its substrates. Based on kinase-catalyzed crosslinking with ATP-photocrosslinking analogs, the K-CLIP method was developed to enrich and identify the kinases of a selected phosphoprotein after immunoprecipitation of the substrate of interest.^[6]^ To modify K-CLIP for discovery of substrates of a selected kinase, a new photocrosslinking analog, ATP-alkyne aryl azide (ATP-AArN_3_) was designed and reported here. Like prior ATP-crosslinkers, ATP-AArN_3_ facilitated covalent linkage of kinase and substrate upon UV irradiation (Figure 3). With the addition of an alkyne handle, immunoprecipitation of the kinase-conjugated substrates was coupled with secondary alkyne-azide click-mediated enrichment to selectively elute bound phosphoprotein substrates, which avoided complicating upstream kinases or kinase-associated proteins (Figure 2). As expected, PDK1, a known kinase that phosphorylates PKA, was not seen in the proteomics list despite being abundant in HEK293.^[7]^ Additionally, PKA regulatory subunits, which bind and inactivate PKA, were seen in the proteomics list, but not enriched (Figure 4B). The absence of PKD1 and PKA regulatory subunits among the hit list confirmed that kinase-focused K-CLIP avoided upstream kinases and associated proteins.

Kinase-focused K-CLIP with ATP-AArN_3_ yielded 5 hits that included two known PKA substrates, CTPS1 and NEK10,^[14, 15]^ which further established the value of the method. CTPS1 generates cytidine triphosphate (CTP), which is necessary for nucleic acid synthesis.^[22]^ PKA phosphorylates CTPS1 in yeast, activating CTP synthase activity.^[23]^ Interestingly, in humans the reverse occurs, with PKA phosphorylation inhibiting CTPS1 synthetase activity, which provides a mechanistic explanation for the influence of PKA activity on cell growth.^[15]^ The Never in mitosis A-related Kinase (NEK) family is involved in ciliogenesis, with NEK10 also required for DNA damage response.^[24]^ Because the kinase-focused K-CLIP method was designed to avoid upstream kinases, the fact that the only kinase observed in the hit list is a known PKA substrate is encouraging. NEK10 interacts with and is phosphorylated by PKA,^[14]^ which controls NEK10 degradation and ultimately ciliogenesis. Because primary cilia act as both tumor suppressors and activators in cancer cells,^[25]^ proper PKA activity is essential in maintaining NEK10 activity and proper ciliogenesis.

Among the three potential PKA substrates identified by K-CLIP, SMC3 was chosen for validation. SMC3 forms a heterodimer with SMC1, and the SMC1-SMC3 complex regulates sister chromatid cohesion.^[26]^ SMC1 and SMC3 are also involved in DNA double strand break (DSB) repair, becoming phosphorylated in response to DNA damage.^[18, 27]^ Similarly, PKA is tied to DNA damage response.^[28]^ In addition, SMC1 is phosphorylated by the ATM kinase, which is activated by the cAMP-PKA signaling pathway.^[29]^ SMC3 also is involved in the cAMP-PKA pathway by modulating the levels of various cAMP-PKA pathway related proteins upon overexpression.^[30]^ Although to our knowledge no kinase of SMC3 has been reported, despite being a known phosphoprotein, PKA and SMC3 are functionally linked. Here we provide the first evidence that SMC3 is a PKA substrate using a cell-based assay (Figure 5).

Methyltransferase-like protein 7A (METTL7A) and Protein sel-1 homolog 3 (SEL1L3) were also discovered as putative PKA substrates using K-CLIP. METTL7A is a thiol methyltransferase (TMT) that methylates thiol-containing drugs.^[31]^ Phosphorylation influences the cellular localization of METTL7A by transport to lipid droplets (LDs),^[16]^ and PKA affects LD dynamics through phosphorylation,^[32]^ suggesting a relationship between METTL7A and PKA. SEL1L3 is a known phosphoprotein, although little is known about its cellular role.^[17]^ Elevated transcription of the SEL1L3 gene is correlated with breast cancer patient survival,^[33]^ and PKA phosphorylates estrogen receptor α to influence breast cancer.^[34]^ The discovery of METTL7A and SEL1L3 as putative substrates highlights the ability for K-CLIP to discover unanticipated phosphorylation-related biology.

Several methods are available for discovery of kinase substrates. Phosphoproteomics coupled with kinase knockdown has been used to identify substrates,^[35]^ although the activity of other similar kinases can alter the substrate profile. Selective inhibitors are also available for many, but not all, kinases and have been used to discovery substrates with phosphoproteomics.^[36]^ Allele-sensitive kinase mutants utilizing base-modified ATP analogs for substrate identification are also available for roughly 10% of human kinases.^[37]^ With the flexibility to identify the substrates of any selected kinase in any complex cellular mixture, kinase-focused K-CLIP offers a useful alternative to available methods. Given prior work identifying the kinases of a selected phosphoprotein substrate,^[6]^ K-CLIP offers a flexible and powerful tool to study any kinase or phosphoprotein substrate of interest from any complex cellular mixture.

Although powerful, a drawback of K-CLIP in this proof-of-concept study was reliance on transient overexpression of a Flag-tagged PKA fusion protein to facilitate mild Flag peptide elution after immunoprecipitation. In addition, overexpression could have created super-physiological concentrations of PKA. As an alternative to overexpression and Flag peptide elution, the kinase-substrate complexes can be immunoprecipitated with a kinase-selective antibody, which are commercially available for many kinases. In this case, elution with sodium dodecyl sulfate (SDS) denaturant, which is compatible with click reaction conditions,^[38]^ can be used before click-mediated secondary enrichment. Another weakness of K-CLIP using ATP-AArN_3_ is the reliance on UV irradiation. Although activation of SMC3 to repair DNA DSB is only reported with ionizing radiation,^[19]^ UV irradiation could have affected the biology of the lysate. To avoid UV light, affinity-based ATP analogs relying on electrophilic crosslinking have been reported,^[39]^ which could be synthesized with an alkyne handle to facilitate the dual enrichment of kinase-focused K-CLIP.

## Conclusion

Kinase-focused K-CLIP was developed using the novel ATP analog ATP-AArN_3_ and used to identify the substrates of PKA. K-CLIP identified SMC3 as a substrate of PKA, which provides a basis for future studies to examine the role of PKA in chromatin cohesion and DNA damage response. Importantly, the study validated K-CLIP as a flexible tool to detect kinase-substrate relationships, whether focused on a target kinase or substrate, which will significantly enhance studies on phosphorylation-mediated cell biology.

## Author Contribution

Hannah J. Bremer: Conceptualization, Investigation, Writing; Mary Kay H. Pflum: Conceptualization, Writing, Supervision, Funding Acquisition

## Data Availability

The data supporting this article have been included as part of the Supplementary Information.

## Supporting information

Supplemental Proteomics Data

## Supplemental Information

Document S1. Materials, instruments, synthetic procedures, Figures S1-S13 Table S1. Excel file containing Proteomics Data, related to Table 1 and Figure 4

## Acknowledgements

We thank the National Institutes of Health (R35GM131821 to M. Pflum, R01GM098285 to Wayne State University Lumigen Instrument Center for High Resolution Mass Spectrometry, and P30ES020957, P30CA022453 and S10OD030484 to Wayne State University Proteomics Core Facility) and Wayne State University for funding, and E. Davis, A. Herppich, and T. Oyewumi for comments on the manuscript. The content is solely the responsibility of the authors and does not necessarily represent the official views of the National Institutes of Health.

## METHODS

### Kinase-catalyzed crosslinking with ATP-AArN_3_

Kinase-catalyzed crosslinking reactions were performed with recombinant PKA kinase (200 ng) in kinase buffer (50 mM HEPES pH 7.6, 150 mM NaCl, 10 mM KCl, 10 mM MgCl_2_). ATP-AArN_3_ **1b** (10 mM final concentration) was added to initiate the reaction (20 μL final reaction volume). Reactions were incubated for 2 hr at 31°C with 300 rpm shaking uncapped under a UV lamp (365 nm), as previously described.^[40]^ Control samples without UV and ATP-AArN_3_ (water used as vehicle) were capped, covered in foil, and incubated in the same incubator. After incubation, 4X Laemmli buffer (5 μL; 277.8 mM Tris pH=6.8, 44.4% (v/v) glycerol, 4.4% LDS, 0.02% bromophenol blue, 10% 2-mercaptoethanol) was added to each sample for a final volume of 25 μL. Reactions were heat denatured for 1 min at 95°C in a sand bath, and proteins were separated using 10% SDS-PAGE. Proteins were then transferred to a PVDF membrane (Immobion P^SQ^, EMD Millipore) for western blot analysis with a PKA primary antibody (mouse, Santa Cruz, cat. No. sc-390548), followed by incubation with secondary antibody (Anti-mouse HRP, Abcam, cat. no. ab97040). Membranes were visualized using a FluoroChem imaging instrument.

### Cell growth and transfection

HEK293 cells (1×10^6^) were grown in T75 flasks with DMEM media supplemented with 10% fetal bovine serum (FBS) and 1% antibiotic/antimycotic at 37°C and 5% CO_2_ atmosphere to 70% confluency. For validation experiments, after a 24 hr growth period, cells were treated with H89 (30 μM in 2% DMSO) or vehicle control (2% DMSO) for 24 hours and then harvested. For K-CLIP studies, Jetprime reagent was used to transfect cells with PKAα-FLAG mammalian expression plasmid DNA (5 μg, Addgene cat. no. 185380) or without plasmid according to manufacturer’s instructions. After transfection, cells were allowed to recover for 36-48 hours to reach confluency before harvesting through centrifugation (1000 rpm, 5 minutes, 4°C) and washing once with Dulbecco’s phosphate-buffered saline (1 mL, DPBS, 10 mM Na_2_HPO_4_, 1.8 mM KH_2_PO_4_, 137 mM NaCl, 2.7 mM KCl, pH 7.4). The cell pellet collected after centrifugation (1000 rpm, 5 minutes, room temperature) was lysed immediately or stored at -80°C. For lysis, cell pellets were incubated with HEPES lysis buffer (500μL; 50mM HEPES, pH 8.0, 150 mM NaCl, 50 mM KCl, 10 mM MgCl_2_, 10% glycerol, 0.5% Triton X-100) supplemented with 1X Xpert protease inhibitor cocktail at 4°C for 40 min with rotation. The supernatant was collected by centrifugation (13,200 rpm, 15 min, 4°C). Protein concentration was determined using the Bradford assay. Lysates were aliquoted for single use and stored at -80°C until use.

### Kinase-focused K-CLIP

Kinase-catalyzed crosslinking reactions were performed with HEK293 lysates containing overexpressed PKA-Flag (500 μg) in kinase buffer (50 mM HEPES pH 7.6, 150 mM NaCl, 10 mM KCl, 10 mM MgCl_2_). ATP-AArN_3_ **1b** (10 mM final concentration) was added to initiate the reaction (75 μL final reaction volume). Reactions were incubated for 2 hrs at 31°C with 300 rpm shaking. The experimental reaction was incubated uncapped under a UV lamp (365 nm), as previously described.^[40]^ Control reactions without UV activation and ATP-AArN_3_ (water as the vehicle) were capped and covered in foil in the same incubator. Flag M2 conjugated beads (30 μL bead slurry) were washed with Tris Buffered Saline (TBS, 500 μL, 20 mM Tris-HCl, 150 mM NaCl, pH 8), added to kinase reactions, and rocked for 3 hours at 4°C. After immunoprecipitation, beads were washed three times with HEPES lysis buffer (500μL) with centrifugation (5,000 rcf for 1 min) to collect the beads. Crosslinked complexes were then eluted from the beads using 3x Flag peptide (40 μL; 0.25 mg/mL in TBS) with 600 rpm shaking for 30 minutes at 4°C. The eluent was collected and added to pre-washed azide agarose resin (80 μL slurry resuspended in 250 μL TBS after one washing with 500 μL of TBS). Click reagents were added (100 μL final volume; 1 mM CuSO_4_, 0.102 mM Tris(benzyltriazolylmethyl)amine (TBTA), and 10 mM TCEP) and incubated for 1 hr at 25°C with shaking while protected from light. After incubation, the bound resin was washed ten times with phosphate binding buffer (PBB, 400 μL; 0.1 M phosphate, pH 7.2, 0.15 M NaCl) and five times with water (400 μL). Proteins were then eluted using acidic elution buffer (300 μL; 0.2% trifluoro acetic acid (TFA), 0.1% formic acid, 80% acetonitrile, 20% water). Eluent was collected and dried using a SpeedVac concentrator. Dried proteins were then resuspended in HEPES lysis buffer (40 μL), 4X Laemmli buffer (10 μL) was added to each sample for a final volume of 50 μL, and samples were heat denatured for 1 min at 95°C. Samples for LC-MS/MS analysis were processed as described below. For western blot analysis, proteins in the reactions were separated using 10% SDS-PAGE, electrotransferred to a PVDF membrane (Immobion P^SQ^), and visualized by incubating with a primary antibody for Flag (Sigma, mouse, cat. # F3165) followed by a secondary antibody (Anti-mouse HRP, Abcam, cat. no. ab97040). Membranes were visualized using a FluoroChem imaging instrument.

### LC-MS/MS sample preparation and in-gel digestion

Kinase K-CLIP for proteomics analysis included three samples that all included transfected HEK293 lysates containing overexpressed PKA-FLAG: a nonspecific bead binding control without ATP-AArN_3_ and UV, an uncrosslinked negative control with ATP-AArN_3_ but without UV, and crosslinking reactions with both ATP-AArN_3_ and UV. Two independent trials were conducted, similar to prior proteomics discovery studies.^[41]^ In-gel tryptic digestion was performed by following a previously published protocol.^20^ After K-CLIP reaction, as described above, proteins were loaded onto 10 % SDS-PAGE gels and only run until proteins had moved one centimeter into the gel. Proteins on gel were visualized with Sypro Ruby total protein stain and an FBTIV-88 ultraviolet trans-illuminator. All proteins in each gel lane were separately excised and cut into 1 mm cubes. Gel pieces were washed with 50 mM NH_4_HCO_3_ (200 μL; 197 mg NH_4_HCO_3_ in 50 mL HPLC water) for 10 min. After removal of the wash, gel pieces were dehydrated with HPLC acetonitrile (250 μL) for 15 min. The liquid was removed, and more acetonitrile (250 μL) was added and removed after 5 minutes. The washing and dehydration steps were then repeated a second time. After washing, samples were dried using a speedvac concentrator (SDP131DDA, Thermo Scientific Savant) for 10 min. Once dry, gel pieces were incubated in 50 mM TCEP/NH_4_HCO_3_ reducing buffer (250 μL; 14.33mg TCEP, 500 μL HPLC water, 500 μL 50 mM NH_4_HCO_3_) for 20 min at 37°C. The buffer was removed, and an alkylation buffer was added (250 μL; 14 mg iodoacetamide, 420 μL HPLC water, 280 μL 50 mM NH_4_HCO_3_). Samples were covered in foil and incubated for 1 hr while rocking at room temperature. The alkylation buffer was removed, and gel pieces were washed and dehydrated twice, as described above. Proteomics grade trypsin (20 μg/mL) was placed on ice and dissolved in 50 mM HCl (100 μL). Gel digestion buffer (900 μL; 800 μL 50 mM NH_4_HCO_3_, 90 μL acetonitrile, 110 μL HPLC water) was added to the trypsin solutions. The gel digestion buffer/trypsin mixture was added to gel pieces (300 μL, each sample), incubated on ice for 15 min, and then incubated for 16 hr at 37°C. After overnight incubation, the digestion solutions were collected and placed on ice. Extraction buffer (200 μL; 500 μL acetonitrile, 500 μL HPLC water, and 2 μL analytical grade formic acid) was added to the gel pieces and sonicated for 15 min. The elution buffer was combined with the digestion solution, dried via speed vac, and stored at -20°C until analysis at Wayne State Proteomics Core Facility.

### LC-MS/MS and ProteomeDiscoverer

Dried samples were resuspended in a solution of 5% acetonitrile, 0.1% formic acid, and 0.005% TFA (15 μL). Each sample (7.5 μL) was separated by reverse phase ultra high-pressure liquid chromatograhy (RP-UHPLC) with an EASY-nLC 100 liquid chromatograph (Thermo Fisher) and C18 columns. An Orbitrap Eclipse Tribrid (Thermo Fisher) mass spectrometer was used to analyzed separated peptides using the following conditions. MS1 spectra were scanned from 375-1600 m/z at 240,000 resolution. MS2 scans were analyzed in the ion trap and peptides were fragmented by collision-induced dissociation (CID). For data analysis, the human protein database from UniProt was used. Searches allowed 2 missed tryptic cleavages. The iodoacetamide derivative of cysteine was set as a fixed modification, while oxidation of methionine and acetylation of peptide N-termini were set as variable modifications. Mass tolerances for parent ions were 20 ppm for the first search, 4.5 ppm for the second search, and 20 ppm for fragment ions. Minimum protein and peptide identification probabilities were specified at 1% false discovery rate (FDR) as determined by a reversed database search, and proteins required 1 unique peptide. All other parameters were used at their default settings. Label free quantification was performed using ProteomeDiscoverer. ProteomeDiscoverer intensities are shown in Table S1A.

To identify enriched substrate proteins in the crosslinking samples, the following calculations were performed. To identify proteins nonspecifically bound to the bead, proteins with higher intensity in the bead binding negative control samples (no UV and ATP-AArN_3_) compared to the crosslinked samples (with UV and ATP-AArN_3_) within each trial were removed (Figure 4A). Fold enrichment values were calculated by dividing the intensities of proteins in crosslinked samples (with UV and ATP-AArN_3_) by the intensities of that same protein in uncrosslinked samples (no UV control) within each trial. Protein hits were expected to give greater enrichment (>1.5-fold) in crosslinking samples compared to uncrosslinked samples in both independent trial (Table S1B and Figure 4A). The final 5 hits were found only in crosslinked, but not uncrosslinked, samples in both independent trials (Table S1B and Table 1).

### Synthesis of ATP-biotin

ATP-biotin was synthesized as previously reported.^[42]^

### In-cell assays

Lysates (1 mg total protein) from HEK293 cells that were treated without (2% DMSO) or with PKA inhibitor H89 (30 μM in 2% DMSO) were incubated without (water used as vehicle) and with ATP-biotin (5 mM) at 31°C for 2 hours with shaking (300 rpm) at 75 μL total volume. After the biotinylation reaction, a small portion of the lysates (100 μg or 10%) were saved as an input control. The remaining samples were diluted with PBB (up to 400 μL total volume), and excess ATP-biotin was removed by filtering with 3 kDa centriprep columns (EMD Millipore, UFC500396) at 14,200 rcf for 30 minutes twice, adding PBB (up to 400 μL total volume) for the second filer step. After filtration, samples were collected, diluted with PBB (up to 400 μL total volume) and incubated with NeutrAvidin resin beads (250 μL bead slurry, Genscript; prewashed once with 500 μL of PBB) for 1 hour at room temperature with rotation. After incubation, the resin was washed ten times with PBB (500 μL) and five times with water (500 μL) to remove nonspecifically bound proteins. Avidin bound biotinylated proteins were eluted by boiling the resin in 2% SDS in water (200 μL) for 7 minutes at 95°C. The eluate was concentrated using a SpeedVac (Thermo scientific), resuspended in 4X Laemmli sample buffer (20 μL), denatured at 95°C for 1 minute, and separated using 12% SDS-PAGE. Proteins were electrotransferred to a PVDF membrane for western blot analysis with SMC3 primary antibody (Cell Signaling, 5696S) and anti-rabbit secondary antibody (Cell Signaling, 7074S). Membranes were visualized using a FluoroChem imaging instrument. Bands were quantified using ImageJ software. Intensity values of SMC3 bands were divided by the SMC3 band intensity in the ATP-biotin reaction sample (set to 100%) and multiplied by 100 to generate a percentage. One-way student t-test was performed using GraphPad Prism version 8.0.

