## Supplemental Data for "Chemoproteomic Profiling of PKA Substrates with Kinase-catalyzed Crosslinking and Immunoprecipitation (K-CLIP)"

**Table of Contents** **Page**

Materials S2

Instruments S2

Synthetic Procedures S2-S4

Figure S1: $^{1}H$ NMR of **4** S5

Figure S2: $^{13}C$ NMR of **4** S5

Figure S3: Mass spectrum of **4** S6

Figure S4: $^{1}H$ NMR of **5** S6

Figure S5: $^{13}C$ NMR of **5** S7

Figure S6: Mass spectrum of **5** S7

Figure S7: $^{1}H$ NMR of ATP-AAr$N_3$ **1b** S8

Figure S8: $^{13}C$ NMR of ATP-AAr$N_3$ **1b** S8

Figure S9: $^{31}P$ NMR of ATP-AAr$N_3$ **1b** S9

Figure S10: Mass spectrum of ATP-AAr$N_3$ **1b** S9

Figure S11: Kinase-catalyzed crosslinking with ATP-AAr$N_3$ **1b** S10

Figure S12: Kinase-focused K-CLIP with Flag-PKA S11

Figure S13: PKA dependent enrichment of SMC3 S12

References S13

### Materials

Fmoc propargyl glycine, 2,2-ethylenedioxybis-(ethylamine), anti-Flag antibody (mouse, cat. # F3165), Flag M2 conjugated beads, copper sulfate, TCEP (tris(2-carboxyethyl)phosphine), proteomics grade trypsin, azide agarose, and 4-aminobenzoic acid were purchased from **Sigma Aldrich**. $Boc_2O$ (di-*tert*-butyl bicarbonate), NMI (N-methylimidazole), and EDCI (1-ethyl-3-(3-dimethylaminopropyl)carbodiimide) were purchased from **Oakwood Chemical**. The disodium salt of ATP and Dulbecco's phosphate-buffered saline (DPBS) were purchased from **Fisher Scientific**. Fmoc-propargylglycine was purchased from **Millipore Sigma**. Bradford reagent and 4X Laemmli sample buffer were purchased from **BioRad**. PKA (cat. # P6000L) was purchased from **New England Biolabs**. PKA selective inhibitor H89 (cat. # B2190) and 3x Flag peptide were purchased from **APExBIO**. PKA antibody (mouse, cat. # sc-390548) wsd purchased from **SantaCruz**. SMC3 primary antibody (cat # 5696S) and secondary goat anti-rabbit HRP antibody (cat. # 7074s) were purchased from **Cell Signaling**. ProQ Diamond Phosphoprotein gel stain, ProQ Diamond Phosphoprotein gel destaining solution,and Sypro Ruby protein stain were obtained from **Invitrogen.** Secondary goat anti-mouse HRP antibody (cat. # ab97040) was purchased from **Abcam**. HEK293 cells was purchased from **ATCC**. FBS (fetal bovine serum) and DMEM media were purchased from **Gibco**. Antibiotic-antimycotic (penicillin/streptomycin/amphotericin B) was obtained from **Hyclone**. The Xpert Protease Inhibitor Cocktail (cat. # P3100-020) was purchased from **GeneDEPOT**. The horse radish peroxidate (HRP) developer for western blotting and Sypro Ruby total protein stain (S12000) was purchased from **ThermoFisher Scientific**. TBTA was obtained from **Click Chemistry.** Immobilon PVDF membrane (Immobion $P^{SQ}$) was obtained from **EMD Millipore**. Jetprime transfection reagent was obtained from **VWR**.

### Instruments

$^1H$ NMR, $^{13}C$ NMR, and $^{31}P$ NMR (Agilent DD2-600 MHz or Agilent MR-400 MHz), along with high-resolution mass spectra (HRMS, Thermo Orbitrap Exploris 120 ESI), were used to characterize novel compounds. The absorbance of ATP analogs was measured by UV-Vis spectrometry (Infinite 200 PRO plate reader, Tecan). A lyophilizer was used to dry the final purified ATP-Alkyne Aryl Azide (Labconoco, FreeZone 4.5). Bradford assay was performed using a fluorimeter (Infinite 200 PRO plate reader, Tecan). Kinase reactions were incubated using a Thermomixer (Multi-therm Heat-shake, Benchmark Scientific). SDS-PAGE and electrotransfer were performed using a transfer cell apparatus (Mini-PROTEAN 3) from BioRad. After Sypro staining, SDS-PAGE gels were visualized using an iBright FL 1500 imaging system (ThermoFisher) using auto visualization setting. After western blot, PVDF membranes were visualized using a FluorChem imager (Protein Simple-Alpha View-FluorChem Q) by scanning for 30 seconds to 4 minutes, or increasing the gain, as needed to visualize crosslinked complexes.

### Synthetic Procedures

Scheme S1. Synthesis of Boc-protected PEG linker

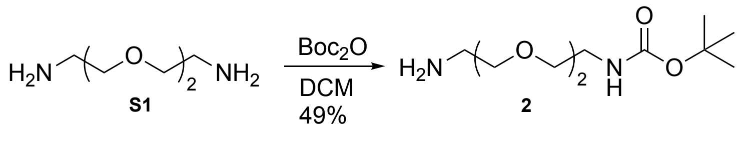


<u>Preparation of Boc-protected PEG linker (*tert*-butyl (2-(2-aminoethoxy)ethyl)carbamate):</u> To a 1000 mL round bottom flask with stir bar was added DCM (350 mL) and **S1** ((2,2-ethylenedioxybis-(ethylamine), 32 mL, 219 mmol, 6 equiv). In a 100 mL beaker with DCM (50 mL) was added di-*tert*-butyl decarbonate ($Boc_2O$, 7.96 g, 36.5 mmol, 1 equiv), which was stirred until fully dissolved. Then, the $Boc_2O$ solution was added dropwise into the solution containing **S1** slowly over 1 hr at room temperature. Once $Boc_2O$ was completely added, the reaction was stirred for 24 hr at room temperature. Production of product was monitored by TLC using 3:1:0.5 EtOH: DCM: $NH_4OH$ with ninhydrin staining (Rf: 0.6). After appearance of the product (roughly 24 hr of reaction), the solvent was evaporated, and the residue was dissolved in water (100 mL). The aqueous layer was then washed with DCM (100 mL) four times, and the combined DCM layers were washed with water (100 mL) four times. The

organic layer was dried with $Na_2SO_4$, filtered, and concentrated to obtain a light-yellow viscous oil (**2**, 4.5g, 49%). Spectral data was consistent with previously reported synthesis.[1-3]

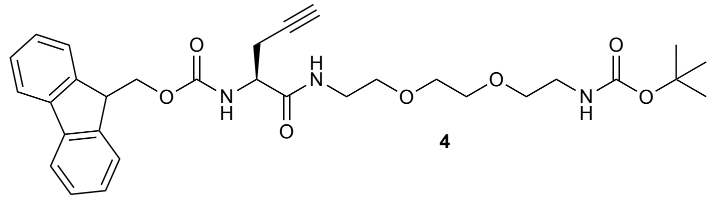

**4**

Preparation of *tert*-butyl (S)-(1-(9H-fluoren-9-yl)-3,6-dioxo-5-(prop-2-yn-1-yl)-2,10,13-trioxa-4,7-diazapentadecan-15-yl)carbamate (**4**): To a 100 mL round bottom flask with a stir bar was added anhydrous THF (5 mL) and Fmoc-propargyl glycine **3** (125 mg, 0.37, 1 equiv). Once dissolved, CDI (91 mg, 0.56 mmol, 1.5 equiv) was added and reaction was stirred for 2 hrs under Ar inert atmosphere at room temperature and monitored by TLC using 3:1 DCM:EtOH to observe the activated amino acid intermediate (Rf: 0.3). Boc-protected PEG linker **2** (178 µL, 0.75 mmol, 2 equiv) was dissolved in THF (2 mL), added to the reaction, and allowed to stir overnight at room temperature. Product formation was monitored by TLC using 8:1 EtOAc:hexanes with ninhydrin staining (Rf: 0.25). Crude product was then purified by column chromatography using 75 mL of silica and 2:1 EtOAc:hexanes (300 mL) and then 100% EtOAc until all product had eluted. The product was collected and dried as a clear oil (**4**, 0.193g, 92%).

$^1$H NMR (400 MHz, cd$_3$od) δ 7.78 (d, J = 7.5 Hz, 2H), 7.66 (d, J = 7.7 Hz, 2H), 7.34 (dt, J = 31.4, 7.4 Hz, 4H), 4.40 (m, 1H), 4.30 (m, 2H), 4.22 (t, 1H), 3.53-3.59 (m, 6H), 3.46 (m, 2H), 3.38 (m, 2H), 3.19 (t, 2H), 2.65-2.58 (m, J = 27.4 Hz, 2H), 2.36 (s, 1H), 1.41 (s, 9H). $^{13}$C NMR (101 MHz, cd$_3$od) δ 208.62, 171.47, 156.82, 143.79, 141.18, 127.41, 126.78, 124.81, 119.54, 79.02, 70.87, 69.88, 69.66, 69.07, 66.74, 53.97, 47.39, 39.81, 39.18, 39.06, 29.27, 27.36, 21.56. ESI calculated $[M+Na]^{+1}$ for $C_{31}H_{39}N_3O_7Na$ 588; Observed 588.

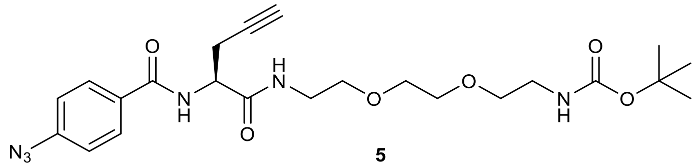

**5**

Preparation of *tert*-butyl (S)-(1-(4-azidophenyl)-1,4-dioxo-3-(prop-2-yn-1-yl)-8,11-dioxa-2,5-diazatridecan-13-yl)carbamate (**5**): To a 100 mL round bottom flask with a stir bar was added **4** (0.193g, 0.34 mmol, 1 equiv) and DCM (4 mL). Once dissolved, piperidine (67 µL, 0.68 mmol, 2 equiv) diluted in DCM (1 mL) was slowly dripped into the flask at room temperature. The reaction was stirred for 15 minutes, and deprotected product was monitored by TLC (8:1 EtOAc:Hexanes with ninhydrin staining, Rf: 0.5). Once starting material was consumed the reaction was diluted with DCM (50 mL) and washed once with sodium bicarbonate (saturated, 50 mL) and water (50 mL). The organic layer was collected and dried. This deprotected amine product of **4** was carried forward to the next step without purification.

To a 100 mL round bottom flask with a stir bar was added 4-azidobenzoic acid (0.112 g, 0.68 mmol, 2 equiv) and DCM (8 mL). EDCI (0.195g, 1.03 mmol, 3 equiv) was separated into three portions, with each portion added every 30 minutes (1.5 hr total) at room temperature. The deprotected amine product of **4** from the prior reaction was separately dissolved in DCM (2 mL) and slowly added to reaction mixture dropwise. The reaction was allowed to stir overnight at room temperature. Product formation was monitored by TLC using 8:1 EtOAc:hexanes with ninhydrin staining (Rf: 0.25). The crude product was purified by column chromatography using 75 mL of silica and 2:1 EtOAc:hexanes, followed by 100% EtOAc. The product was collected as a clear oil (**5**, 0.077g, 46% over two steps). $^1$H NMR (400 MHz, cdcl$_3$) δ 7.83 (d, J = 8.2 Hz, 2H), 7.07 (d, J = 8.6 Hz, 2H), 6.85 (bs, 1H), 5.06 (bs, 1H), 4.75 (bs, 1H), 3.56 (m, J = 25.0 Hz, 12H), 3.28 (m, 2H), 2.86 (m, 1H), 1.82 (s, 1H), 1.43 (s, 9H). $^{13}$C NMR (151 MHz, cdcl$_3$) δ 187.24, 169.52, 166.04, 128.99, 128.74, 119.00, 118.90, 79.44, 71.90, 70.39, 70.25, 70.17, 69.58, 51.89, 40.29, 39.58, 29.67, 28.40, 22.61. ESI calculated $[M+Na]^{+1}$ for $C_{23}H_{32}N_6O_6Na$ 511; Observed 511.

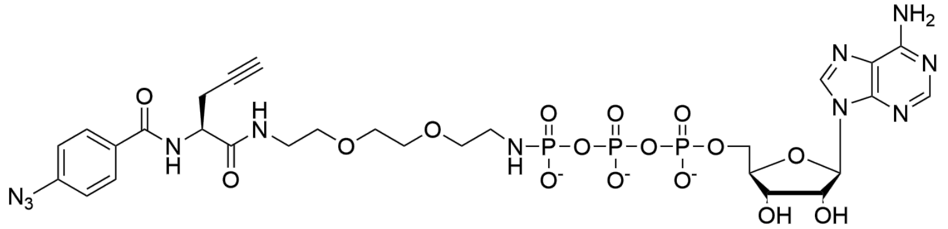


**1b**

Preparation of ATP-Alkyne Aryl Azide (**1b**): To a 100 mL round bottom flask with stir bar was added **5** (0.077g, 0.012 mmol) and 1,4-dioxane (10 mL). The reaction was cooled to 0˚C, followed by addition of HCl (1.5 mL, 4M). The reaction was stirred overnight at room temperature, and deprotected product was monitored by TLC (3:1:0.5 DCM:EtOH:$NH_4OH$, Rf: 0.3). Solvent was then evaporated in vacuo. The resulting deprotected azido propargyl glycine amine crude oil was evaporated with methanol (15 mL) and carried forward to the next step without purification.

To a 50 mL plastic conical tube with stir bar was added the disodium salt of ATP (50 mg, 0.09 mmol, 1 equiv), EDCI (156.4 mg, 0.82 mmol, 9 equiv), and water (1 mL). The reaction was stirred until the solid was dissolved. Then, N-methylinidazole (NMI, 72 µL, 0.9 mmol,10 equiv) was added and the pH was adjusted to 6.2-6.8 with 0.5 M aqueous HCl. Then, the deprotected azido propargyl glycine amine from the prior reaction (153.7 mg, 0.4 mmol, 6 equiv) dissolved in aqueous HCl (1 mL of 0.5M) was added. The pH was then adjusted to 6.2-6.8with 0.5 M aqueous HCl or 0.5 M aqeous NaOH, as needed, and the reaction was stirred for 2 hr at room temperature. Product formation was monitored by TLC (3:1.5:0.5 i-PrOH:$NH_4OH$:$H_2O$, Rf: 0.5). After 2 hrs, the product was isolated by anion exchange chromatography (A-25 Sephadex, 4.5g) with TEAB buffer (triethylammonium bicarbonate buffer; 1 M TEA in water, adjust pH to 8-8.5 by adding dry ice) using a stepwise elution (100 mL of 50 mM, 100 mL of 100 mM, 100 mL of 250 mM, and 200 mL of 370 mM TEAB buffer). The product was collected with 370 mM TEAB fractions, frozen, and lyophilized to yield a white powder (**1b,** 57% yield over two steps) The concentration was measured by absorbance using a UV-Vis spectrophotometer (ε = 15,400 $M^{-1}cm^{-1}$). UV-Vis spectroscopy (100 mM HEPES pH-7.4): 260 $cm^{-1}$.

$^1$H NMR (600 MHz, $D_2O$) δ 8.38 (d, J = 23.1 Hz, 1H), 8.12 (d, J = 32.4 Hz, 1H), 7.63 (dd, J = 32.1, 8.4 Hz, 2H), 6.98 (dd, J = 46.6, 8.3 Hz, 2H), 5.96 (d, J = 35.9 Hz, 1H), 4.48 (t, 1H), 4.40 (d, 1H), 4.23 (s, 1H), 4.08 (s, 2H), 3.44-3.27 (m, 12H), 2.89 (t, 1H), 2.68 (d, 2H), 2.31 (s, 1H). $^{13}$C NMR 191.09, 172.28, 152.59, 129.10, 129.03, 124.26, 118.94, 118.83, 92.33, 86.58, 83.98, 74.19, 71.87, 70.34, 69.51, 69.45, 68.92, 66.30, 65.51, 58.78, 53.13, 42.15, 40.67, 39.26, 10.43, 7.30. $^{31}$P NMR (162 MHz, $d_2o$) δ -1.52 (d, J = 20.5 Hz), -11.53 (d, J = 19.1 Hz), -22.88 (t, J=20.2 Hz). HRMS (ESI-Orbitrap) m/z: $[M-H]^{-1}$ calculated for $C_{28}H_{38}N_{11}O_{16}P_3$ 876.1638; Observed 876.1634.

Resuspension and storage of ATP-AAr$N_3$ **1b** Because all ATP analogs, including ATP-AAr$N_3$ **1b**, are highly prone to hydrolysis and degradation in aqueous storage buffers, care must be taken when resuspending and storing dried ATP analogs for later use. For resuspension, the dried power of ATP-AAr$N_3$ **1b** after lyophilization was mixed to HEPES storage buffer (100 µL, 100 mM HEPES, pH=7.4). To determine the concentration, absorbance was taken with a UV-Vis spectrophotometer, and the concentration was calculated using beers law (A=εbc) where A = the absorbance at 254 nm, ε = 15.4 x $10^3$ $M^{-1}$ $cm^{-1}$, b = path length (cm), and c = concentration of ATP-AAr$N_3$ **1b**. The resuspended solution was distributed into tubes as single use aliquots (typically 5 or 10 µL) and store at -80 °C, which avoids freeze-thaws to prevent degradation. ATP-AAr$N_3$ **1b** can be stored up to 1 year as a dry powder at -80 °C or up to 6-8 months after resuspended in buffer at -80 °C. Purity can be assessed by TLC analysis (silica; 3:1.5:0.5 isopropanol:ammonium hydroxide:water; $R_f$: 0.4), with minimal degradation to ATP (Rf: 0.0) expected.

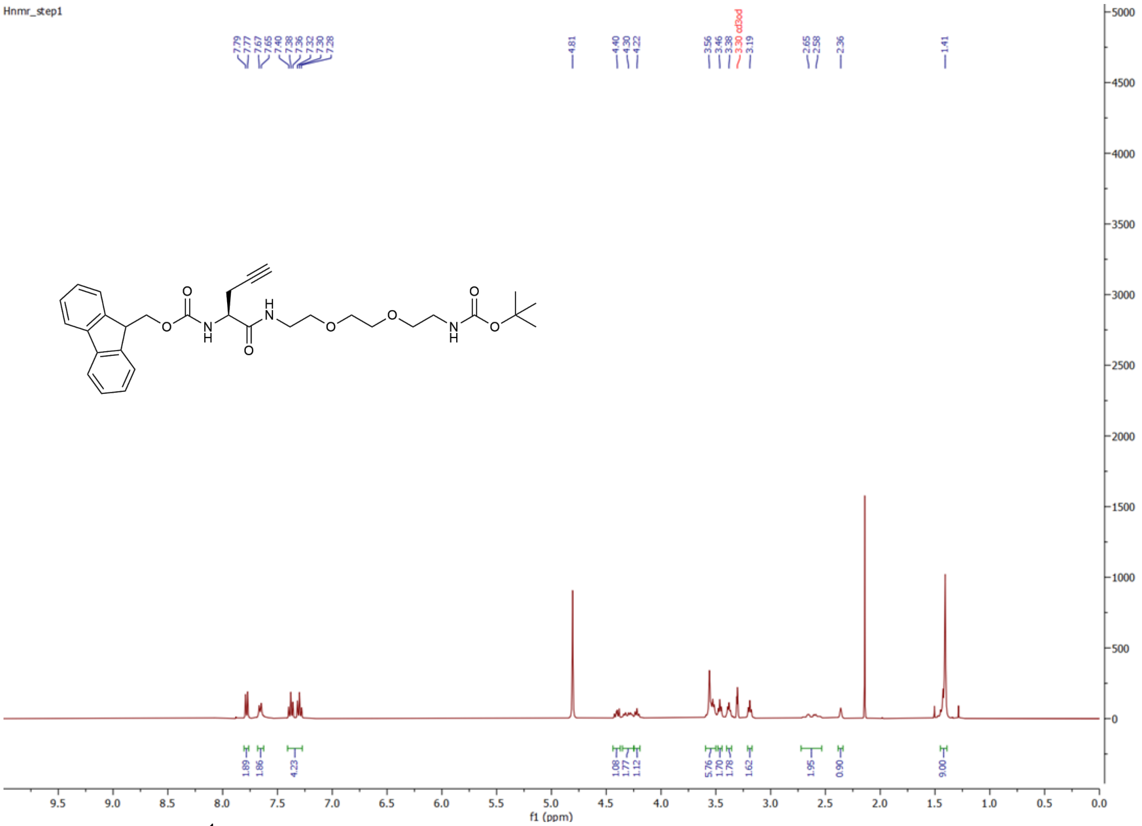

**Figure S1.** $^{1}$H NMR tert-butyl (S)-(1-(9H-fluoren-9-yl)-3,6-dioxo-5-(prop-2-yn-1-yl)-2,10,13-trioxa-4,7-diazapentadecan-15-yl)carbamate **4** recorded in $CD_3OD$. The peak at δ 3.3 corresponds to $CD_3OD$, and the peak at δ 4.8 corresponds to water.

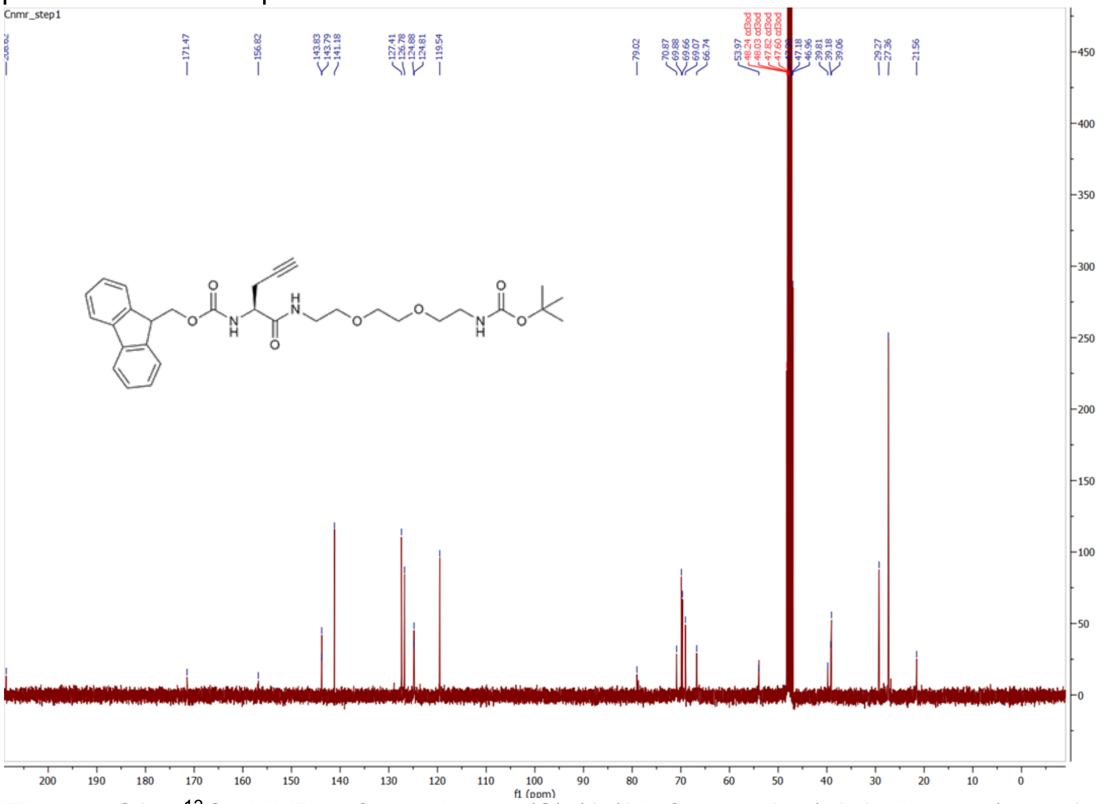

**Figure S2.** $^{13}$C NMR of tert-butyl (S)-(1-(9H-fluoren-9-yl)-3,6-dioxo-5-(prop-2-yn-1-yl)-2,10,13-trioxa-4,7-diazapentadecan-15-yl)carbamate **4** recorded in $CD_3OD$. The peaks at δ 48.24 to 47.60 corresponds to $CD_3OD$.

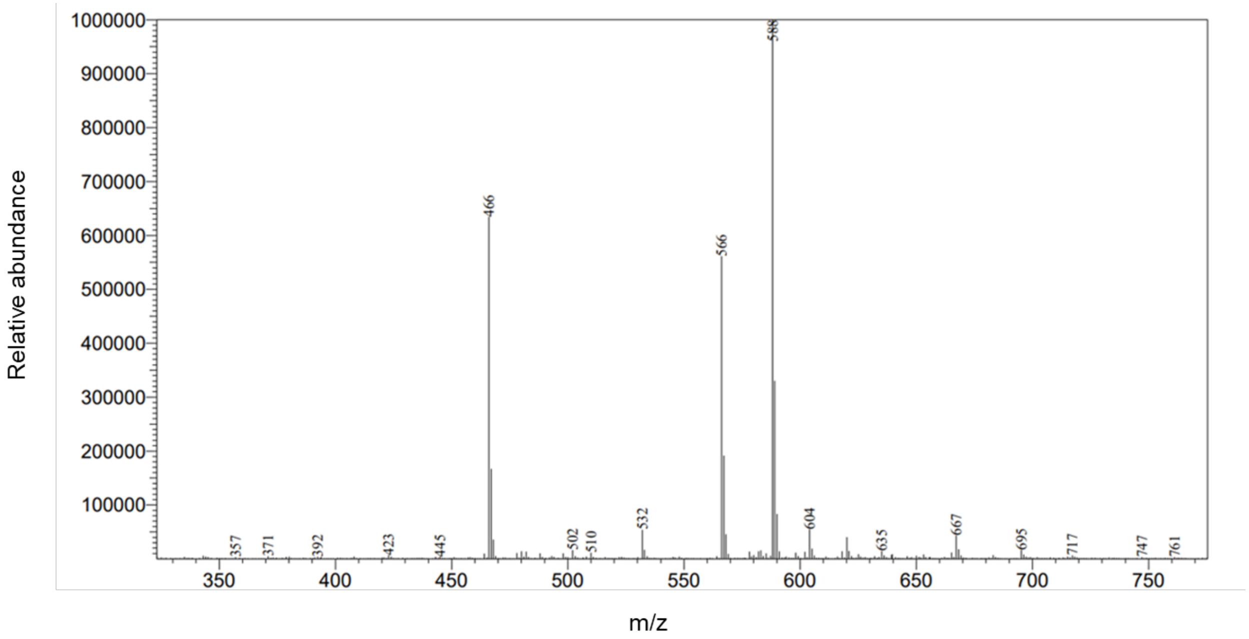


**Figure S3.** Electrospray Ionization (ESI) tert-butyl (S)-(1-(9H-fluoren-9-yl)-3,6-dioxo-5-(prop-2-yn-1-yl)-2,10,13-trioxa-4,7-diazapentadecan-15-yl)carbamate 4. Calculated $[M+Na]^{+1}$ for $C_{31}H_{39}N_3O_7Na$ - 588; Observed – 588.

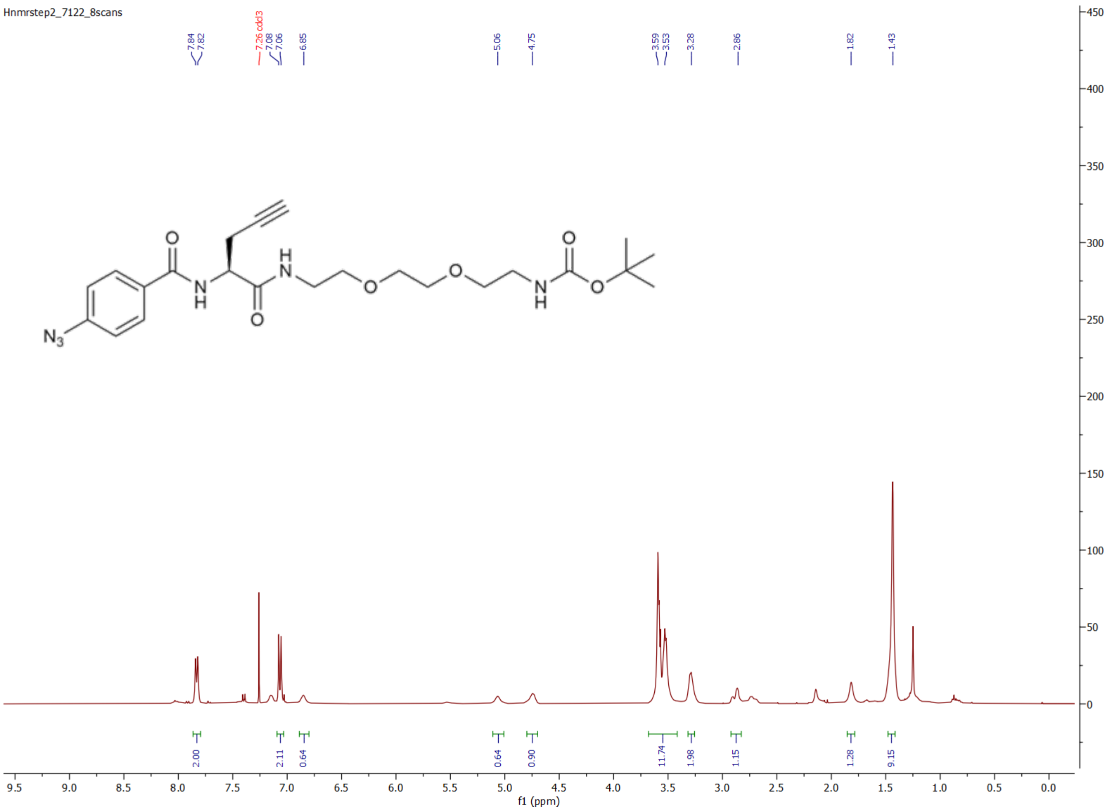


**Figure S4.** $^1$H NMR tert-butyl (1-(4-azidophenyl)-1,4-dioxo-3-(prop-2-yn-1-yl)-8,11-dioxa-2,5-diazatridecan-13-yl)carbamate **5** recorded in $CDCl_3$. The peak at δ 7.26 corresponds to $CDCl_3$. The peak at δ 2.17 corresponds to acetone. The peak at δ 1.26 corresponds to ethyl acetate.

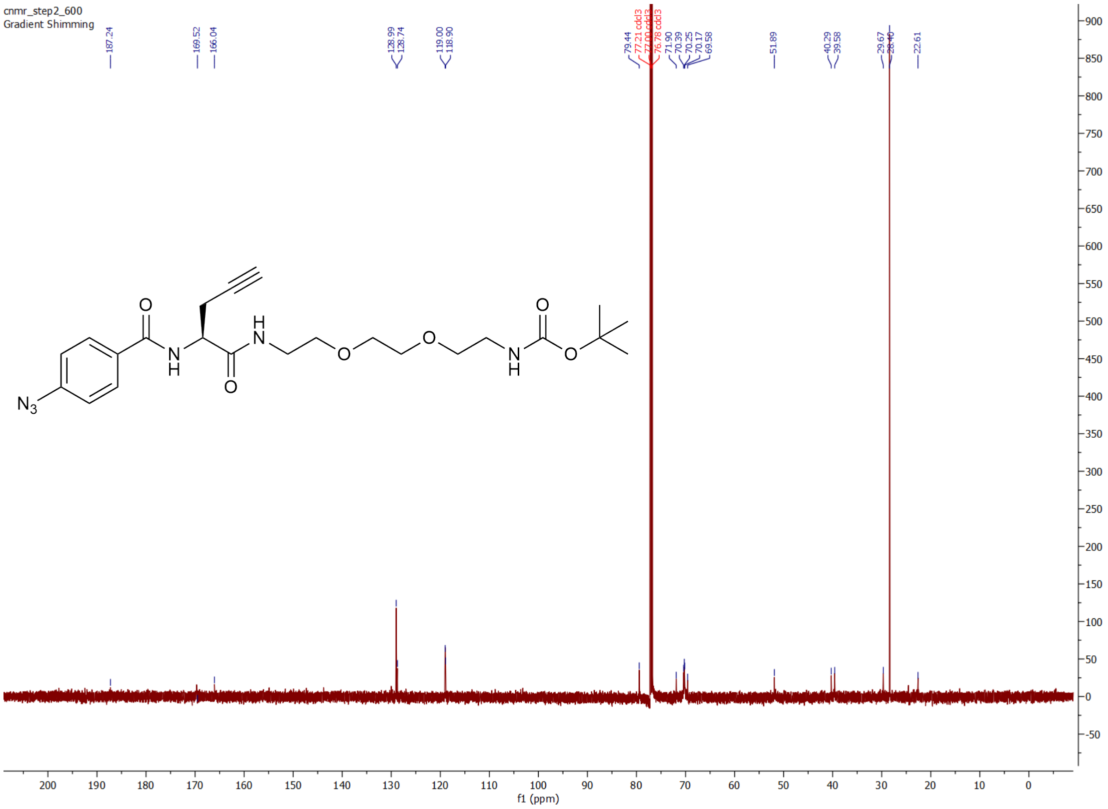


**Figure S5.** $^{13}C$ NMR of tert-butyl (1-(4-azidophenyl)-1,4-dioxo-3-(prop-2-yn-1-yl)-8,11-dioxa-2,5-diazatridecan-13-yl)carbamate **5** recorded in $CDCl_3$. The peaks at δ 77.21 to76.78 corresponds to $CDCl_3$.

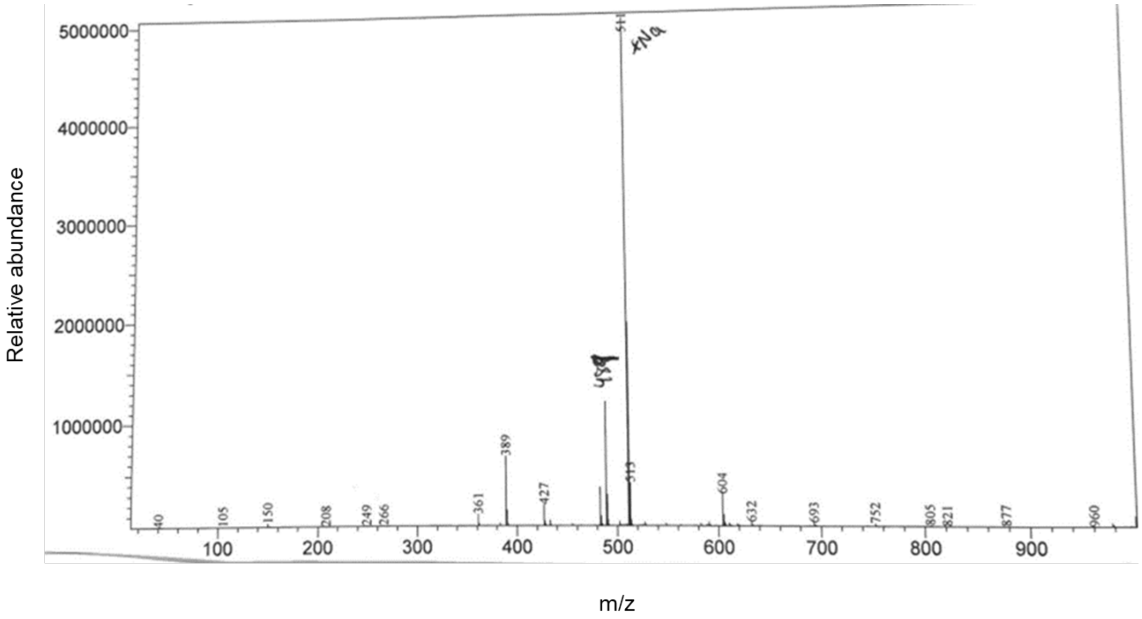


**Figure S6.** Electrospray Ionization (ESI) of tert-butyl (1-(4-azidophenyl)-1,4-dioxo-3-(prop-2-yn-1-yl)-8,11-dioxa-2,5-diazatridecan-13-yl)carbamate **5**. Calculated $[M+Na]^{+1}$ for $C_{23}H_{32}N_6O_6Na$ - 511; Observed - 511.

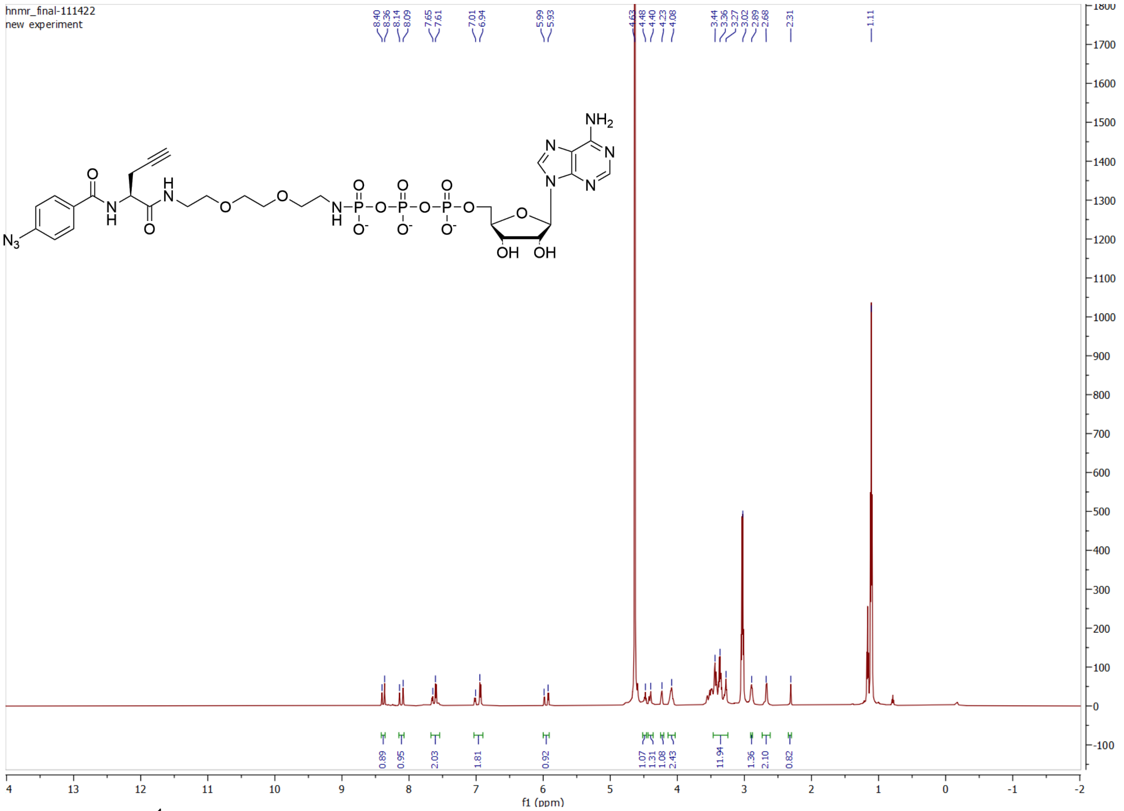

**Figure S7.** $^{1}$H NMR of ATP-AArN$_3$ **1b** recorded in $D_2O$. The peak at δ 4.65 corresponds to $D_2O$, and the peaks at 3.01 and 1.10 correspond to the $CH_3$ and $CH_2$ of the triethylamine counterion salt, respectively.

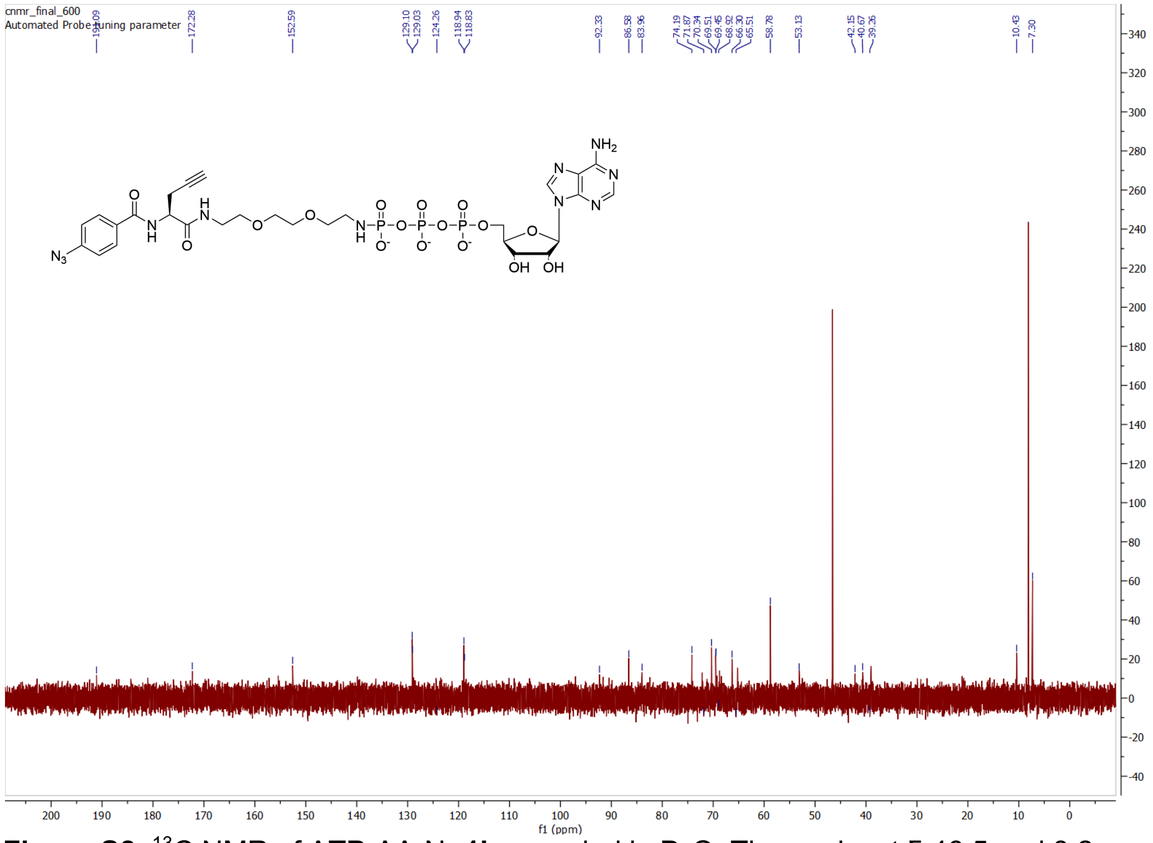

**Figure S8.** $^{13}$C NMR of ATP-AArN$_3$ **1b** recorded in $D_2O$. The peaks at δ 46.5 and 8.2 correspond to the $CH_3$ and $CH_2$ of triethylamine, respectively.

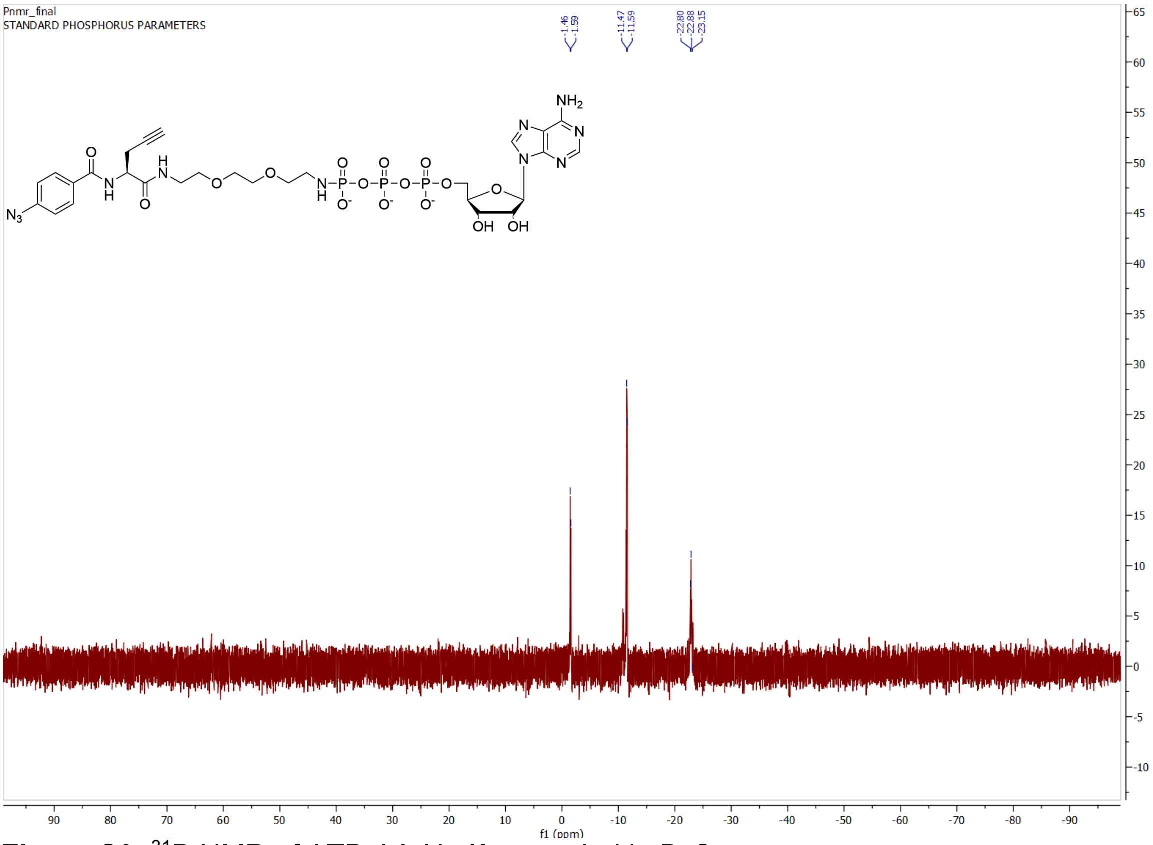


**Figure S9.** $^{31}P$ NMR of ATP-AArN$_3$ **1b** recorded in $D_2O$.

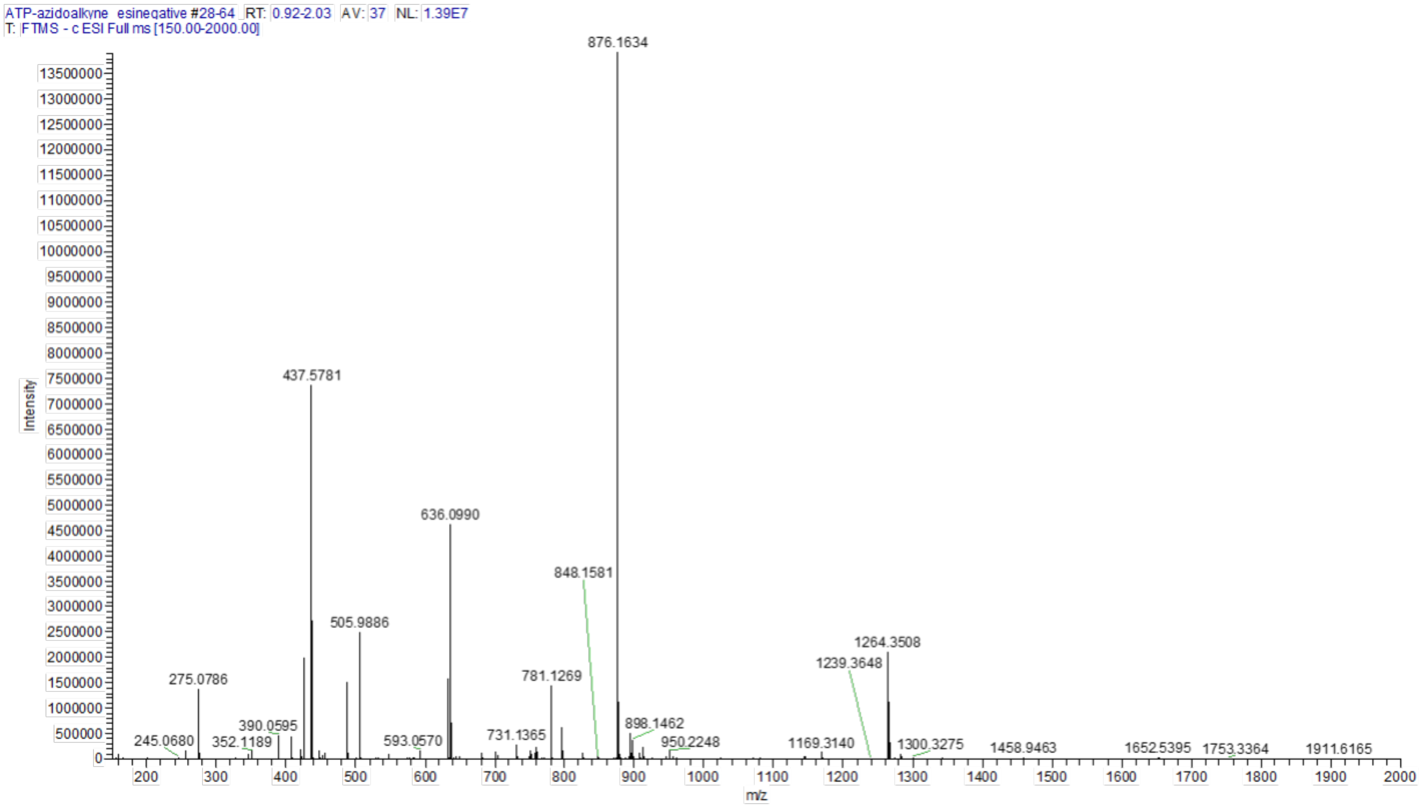


**Figure S10.** Electrospray Ionization (ESI) High resolution mass spectrum (HRMS) of ATP-AArN$_3$ **1b**. Calculated $[M-H]^{-1}$ for $C_{28}H_{38}N_{11}O_{16}P_3$ 876.1638; Observed 876.1634.

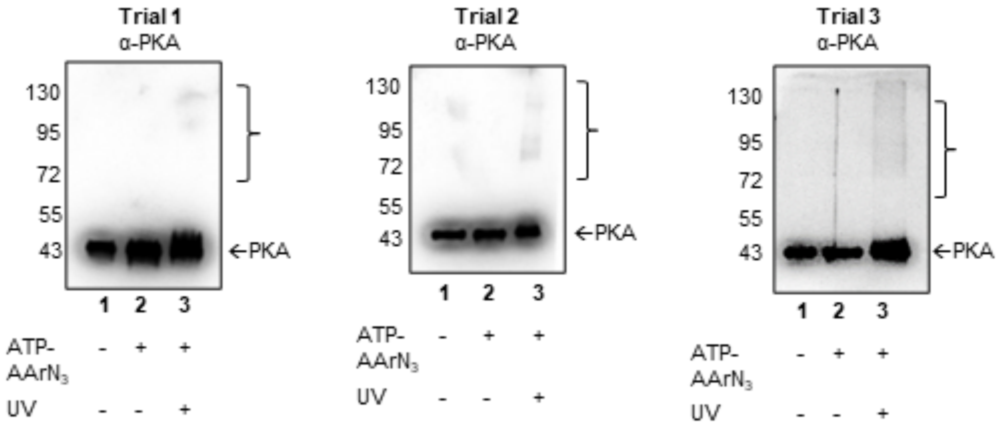


**Figure S11.** Kinase-catalyzed crosslinking with ATP-AArN$_3$ and PKA. Recombinant PKA was incubated without (lane 1) or with ATP-AArN$_3$ (10 mM; lanes 2-3) in the absence (lane 2) or presence (lane 3) of UV light (365 nm) for 2 hours at 31˚C. After crosslinking, proteins were separated via SDS-PAGE and visualized by western blot analysis with a PKA (α-PKA) antibody. Arrows indicates uncrosslinked PKA (43 kDa), and brackets indicate crosslinked PKA dimers (86 kDa) and trimers (129 kDa). Molecular markers are indicated to the left of gel images. Three trials are shown, with a truncated image of trial 2 is shown in Figure 3A.

A.

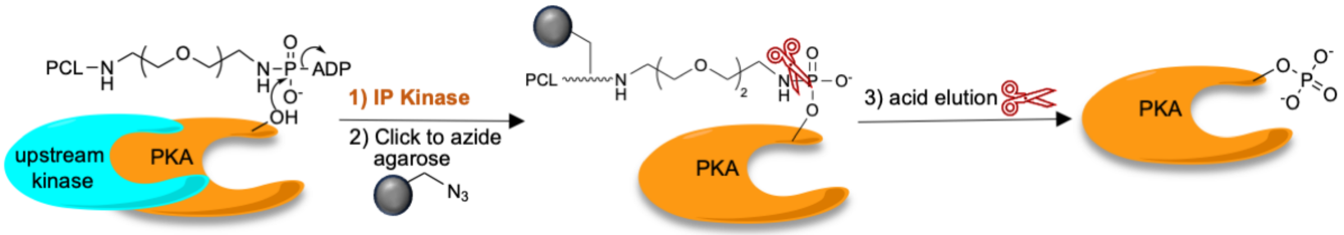


B.

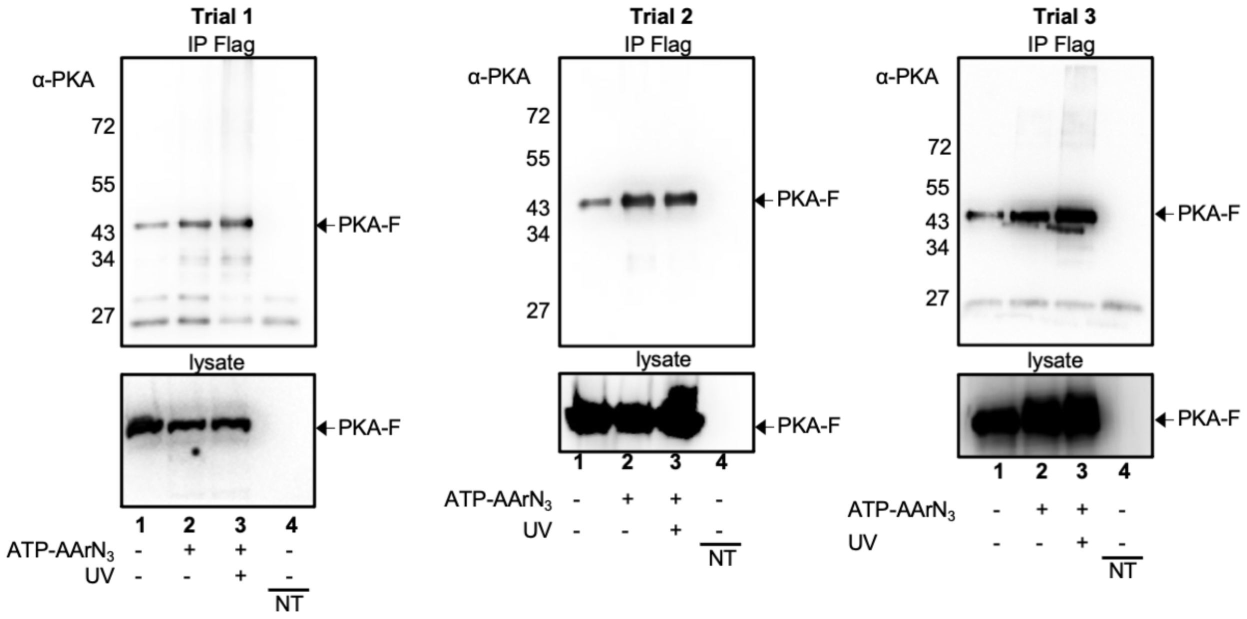


**Figure S12.** PKA-focused K-CLIP with gel analysis. (A) Schematic showing the expected enrichment of PKA using K-CLIP without UV-mediated crosslinking. Cellular lysates containing overexpressed PKA-Flag were incubated with ATP-AArN$_3$ without UV light, which will label PKA as a substrate but not crosslink to other proteins. Labeled PKA was isolated by immunoprecipitation (IP), eluted, and subsequently attached to azide-modified resin via click reaction of the alkyne group within the crosslinker. Acid cleaves the phosphoramidate bond within the crosslinker (scissors), releasing only phosphoproteins from the protein-bound resin, which included PKA. (B) HEK293 cells were transfected without (lane 4, NT = nontransfected) or with (lanes 1-3) a PKA-Flag expression plasmid. The resulting lysates were incubated with (lanes 2 and 3) or without (lanes 1 and 4) ATP-AArN$_3$ (10 mM). PKA crosslinked complexes were isolated by immunoprecipitation via the Flag tag. Enriched proteins were eluted with Flag peptide, and a secondary enrichment with agarose azide resin was performed using click chemistry. Proteins were eluted using acid, separated using SDS-PAGE, and visualized using Flag antibody western blotting. Three trials are shown, with a truncated image of trial 2 shown in Figure 3B.

A.

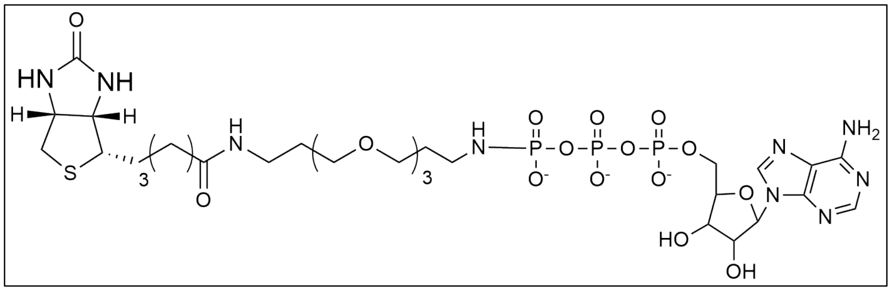

B.

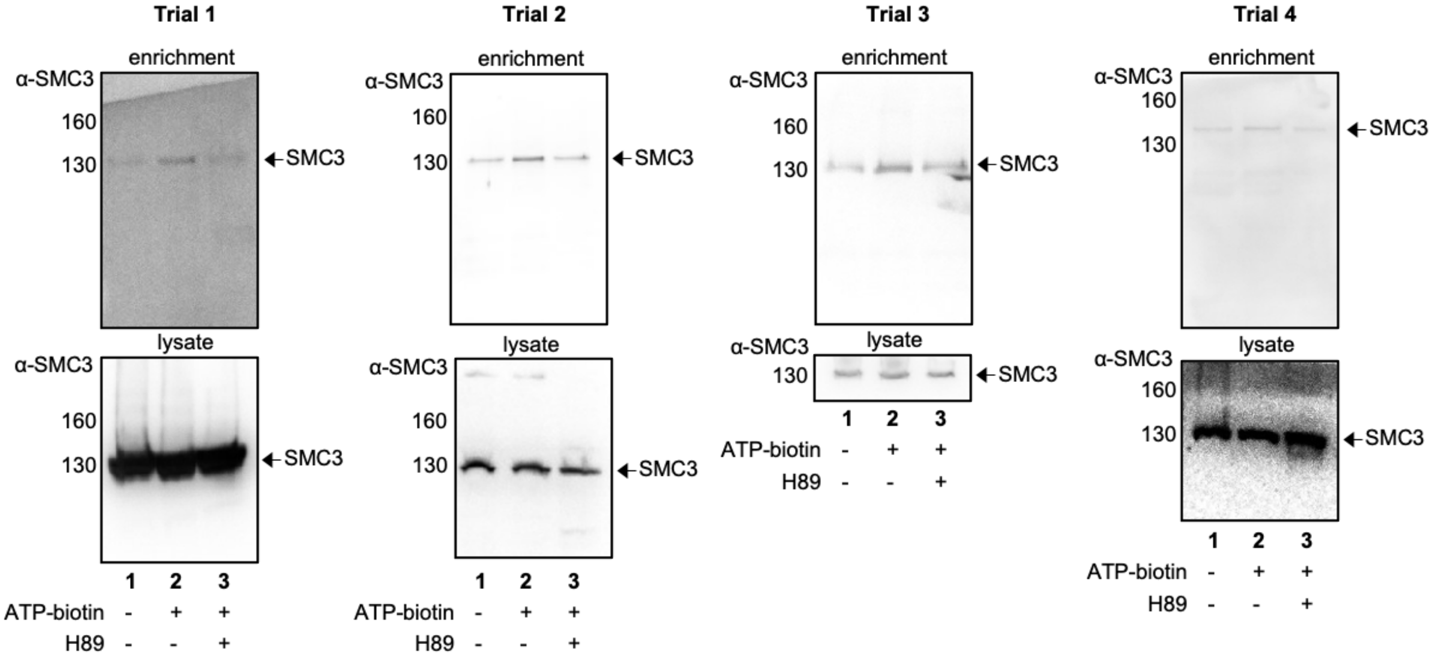


C.

| trial | 1 | 2 | 3 | 4 | mean | SE |
|---|---|---|---|---|---|---|
| bead | 85 | 49 | 44 | 87 | 66 | 23 |
| ATP-biotin | 100 | 100 | 100 | 100 | 100 | |
| H89 | 95 | 57 | 81 | 80 | 78 | 16 |

**Figure S13.** PKA dependent enrichment of SMC3 (A) Structure of ATP-biotin used in kinase-catalyzed biotinylation for the in-cell kinase assays (see Figure 5a of the main manuscript). (B) HEK293 cells were treated without (2% DMSO) or with PKA inhibitor H89 (30 μM in 2% DMSO). The resulting lysates were incubated with ATP-biotin. Biotinylated proteins were enriched with NeutrAvidin resin and separated by 12% SDS-PAGE. Input lysates before enrichment were separated as load controls. SMC3 levels were visualized by immunoblotting with SMC3 specific antibody (α-SMC3). Truncated images of western blot gels from trial 2 are shown in Figure 5B. (C) SMC3 bands from four independent trials in part B were quantified using ImageJ and normalized as a percentage in the ATP-biotin samples (lane 2, set to 100%), with mean and standard error (SE) shown. A histogram with the % of SMC3 biotin enriched from the table is shown in Figure 5C.
